# Resolving differential vascular graft remodeling using longitudinal multiphoton tracking in a 3D culture platform

**DOI:** 10.64898/2026.02.28.708759

**Authors:** David R. Maestas, Trin R. Murphy, Katarina M. Martinet, Tracey Moyston, Leon Xuanyu Min, Ali Behrangzade, Brock J. Pemberton, Jincy Joy, Sang-Ho Ye, George S. Hussey, Mohamad Azhar, William R. Wagner, Jonathan P. Vande Geest

## Abstract

The long-term performance of tissue-engineered scaffolds, particularly small-diameter vascular grafts, is shaped by remodeling events at the tissue-graft interface, yet these processes remain difficult to resolve longitudinally and at microstructural resolution in conventional implantation models. Here we develop an organotypic artery-graft model that preserves cylindrical vessel geometry and enables non-destructive label-free multiphoton monitoring of interface remodeling. Using second harmonic generation and two-photon excited fluorescence, we capture evolving fibrillar collagen architecture and cellularization over time, demonstrate compatibility with multiple biomaterial classes, and show integration with rat and mouse explants, live-cell dyes, and fluorescent reporter tissues. The platform resolved distinct remodeling responses to transforming growth factor-β isoforms (TGF-β1, -β2, and -β3), with differential shifts in collagen-fiber distributions, accompanied by changes in matrix-remodeling and contractile gene expression. Across two graft designs, culture-derived remodeling phenotypes, collagen fiber distributions, and initial trajectories agreed with those observed in long-term 6-month interpositional explants. Together, these results establish an accessible intermediate platform for interrogating artery-graft remodeling, tracking these trajectories, and prioritizing graft designs through interface-resolved outcomes before and alongside animal implantation studies.

## Introduction

Cardiovascular disease remains the leading cause of global mortality, driving the need for small-diameter vascular replacements when autologous conduits are unavailable or insufficient^1-3^. Tissue-engineered vascular grafts (TEVGs) have emerged as promising alternatives, yet their clinical translation remains limited^4-6^. A central challenge is that the early remodeling trajectories that lead to durable integration versus maladaptive fibrosis or stenosis are difficult to resolve longitudinally and non-invasively, even though cellularization and extracellular matrix (ECM) organization at the tissue-graft interface may be among the most informative predictors of long-term patency^4-8^. Although animal implantation studies remain essential for TEVG validation, they are resource-intensive and rely heavily on endpoint analyses and macroscale longitudinal monitoring. As a result, advancing an acellular graft from benchtop fabrication and mechanical characterization to both small-animal and large-animal testing requires substantial time and resources. Reflecting this developmental burden, only a relatively small number of TEVG publications include *in vivo* implantation outcomes, and among those that use acellular grafts, 59% were performed in rodent interpositional models^9-11^. Together, these limitations define a translational bottleneck between benchtop characterization and *in vivo* testing, underscoring the need for intermediate models that can longitudinally assess and map potential tissue-scaffold remodeling dynamics to de-risk graft designs before commitment to resource-intensive animal implantation studies^9-11^.

Model culture systems with strong predictive alignment to *in vivo* outcomes remain difficult to achieve, though recent breakthroughs in three-dimensional (3D) cultures and tissue explant *ex vivo* systems have improved upon the predictive power and biological relevance relative to monolayer cultures^12-14^. However, TEVG-relevant models remain limited, ranging from monolayer cultures that poorly recapitulate the *in vivo* microenvironment, to modern vascular bioreactors that provide tunable luminal flow and axial loading^15-17^. Although these dynamic systems may be essential when hemodynamic conditioning is required, these systems are inherently low-throughput, resource intensive, limited to macro-resolution longitudinal readouts, and rely heavily upon destructive endpoint assays to assess tissue-graft integration states. Notably, the enduring utility of the aortic ring assay, a widely used static explant culture model of angiogenesis, demonstrates that arterial outgrowth and early remodeling can proceed without imposed luminal flow and axial loading, suggesting that these inputs are not strictly required for shorter-term *ex vivo* models. What remains lacking is an accessible intermediate model that preserves cylindrical artery-graft geometry while enabling non-destructive, high-resolution, longitudinal assessments of ECM remodeling and cellularization at the tissue-graft interface, a critical region where remodeling trajectories can diverge toward integration or graft stenosis^18-25^.

In this work, we present a 3D artery-graft interface culture platform that enables longitudinal, quantitative assessment of early remodeling trajectories and cellularization at the tissue-graft interface, supporting graft screening and design evaluation before or alongside animal implantation studies. By directly coupling rodent aortic explants to acellular TEVGs, the platform preserves cylindrical tissue-graft geometry *ex vivo* and, when paired with label-free multiphoton imaging, enables repeated non-destructive monitoring of fibrillar collagen remodeling by second harmonic generation (SHG) alongside endogenous two-photon excited fluorescence (2PEF) signatures arising from cellularization and scaffold-associated signals^26,27^. Using fiber-level quantification of SHG images, we track evolving collagen architecture over culture duration, validate cellular 2PEF signatures using live and fixed nuclear staining, and confirm compatibility with standard gene expression, histology, and immunostaining endpoint assays^28-30^. Across two distinct TEVG designs, we found that culture-derived remodeling phenotypes and collagen microarchitectures recapitulate key features of long-term explant outcomes, supporting the platform’s ability to identify and parallel aspects of remodeling trajectories *in vivo*. To support broader applicability, we demonstrate compatibility with murine and fluorescent reporter models and show that multiple graft biomaterials classes can also be utilized for longitudinal tracking. Lastly, we demonstrate that the platform is sensitive to subtle biochemical perturbations, as treatments with three transforming growth factor-β isoforms (TGF-β1, -β2, -β3) induce distinct shifts in SHG-derived fiber architecture distributions that coincide with complementary changes in collagen gene expression^31,32^.

## Results

### A modular artery-graft *ex vivo* culture to preserve interface geometry for longitudinal imaging

We first engineered a modular organotypic artery-TEVG culture designed to preserve the cylindrical geometry of interpositional graft interfaces while enabling longitudinal imaging from upright or inverted imaging systems (**Fig. 1a-c**). To elevate the constructs above the culture surface and closer to the media-air interface, we used purchasable PTFE-coated mandrels and 3D-printed polycarbonate hexagon-shaped holders (**Fig. 1d**). Rat abdominal aortas were harvested, cleaned of perivascular tissue, mounted on these supports and coupled graft segments in a controlled configuration (**Fig. 1d-g**).

**Fig. 1.**
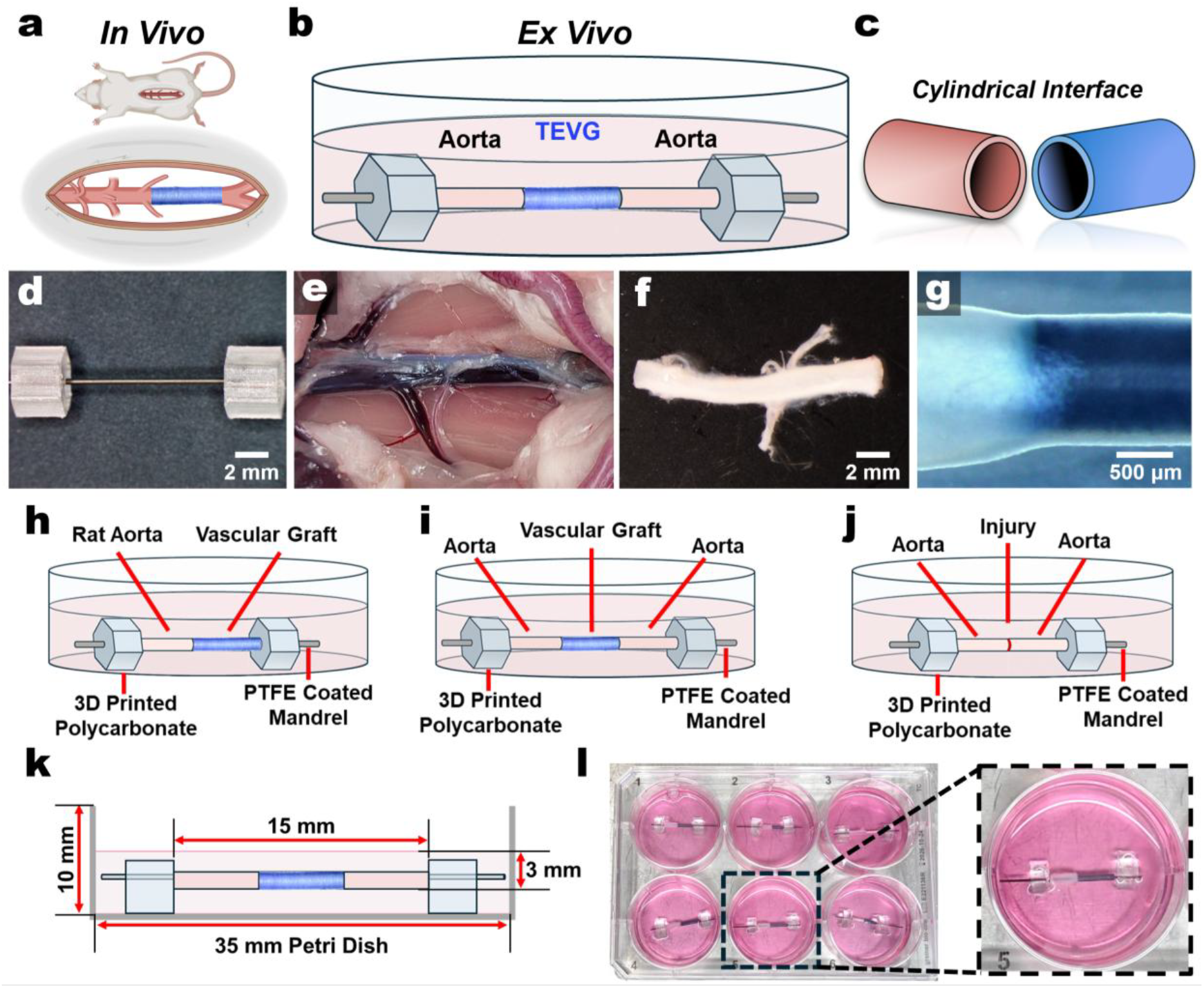
Design of the 3D Artery-Graft Culture Platform. **a-c**. Graphical visualization of rat interpositional grafts, a 3D culture schematic, and the cylindrical geometry interface between them. **d**. Image of assembled parts of the 3D culture before sample additions: a PTFE coated stainless-steel mandrel and 3D-printed polycarbonate hexagon sample holders. **e**,**f**. Isolation and preparation of rat aortas. **g**. Representative zoom-in of the tissue-TEVG interface. **h-j**. Illustrations of platform variations depicting (**h**) a single connection, (**i**) an interpositional (double) connection, and (**j**) a transected and rejoined aorta to model aortic injury and/or autologous grafts. **k**. Schematic showing culture dimensions. **l**. Image of a 6-well plate of single joined cultures.

This modular framework supports three complementary configurations to address distinct questions in graft design: a single artery-graft interface to track cellular or collagen progress from the artery to the graft, an interpositional (double-interface) configuration to model cellularization and remodeling across both anastomotic junctions, and an aortic injury/autologous graft model to compare remodeling and repair when coupled segments are living tissues (**Fig. 1h-j, Extended Data Fig. 1**). We standardized our grafts and aortic segments lengths to ∼5 mm. Across configurations, approximate diameter matching between host vessel and graft improved interface stability, facilitated longitudinal monitoring of cellularization and remodeling, and reduced telescoping at the tissue-graft junctions (**Extended Data Fig. 2, Fig. 1j**). We selected segments lengths so that the assembled constructs fit within 35 mm dishes and 6-well plates to support lower-cost and higher-throughput parallel studies without the need for specialized bioreactor hardware and control systems (**Fig. 1k,l**). Although we treated each aortic explant as a single biological replicate in this work, the modular platform intrinsically supports paired testing of distinct aortic segments from the same vessel under matched or different conditions.

**Fig. 2.**
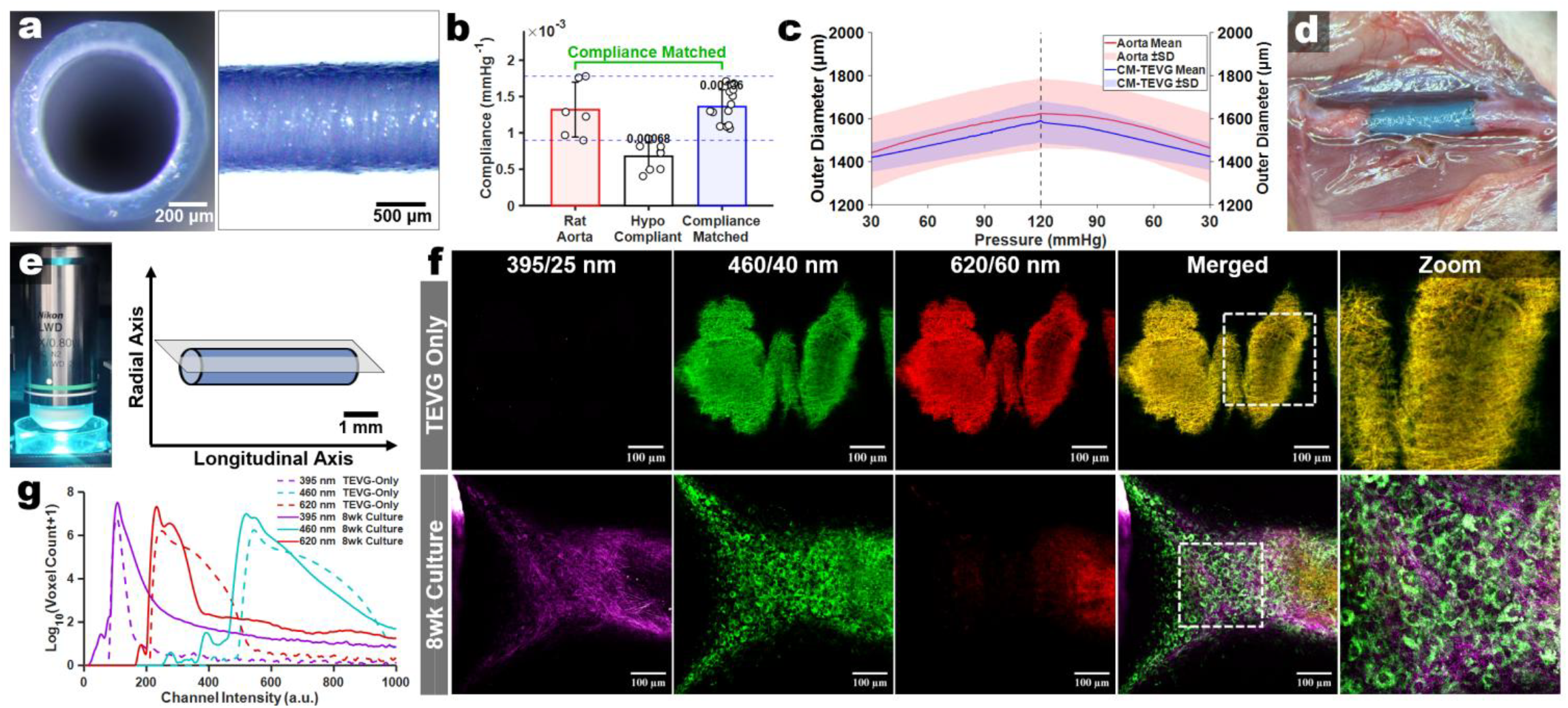
Identifying the spectral signature of our chosen compliance-matched TEVG design. **a**. Representative acellular trilayered TEVG (cross-section and longitudinal view). **b**. Circumferential compliance of native rat abdominal aorta, a hypocompliant TEVG (thickened middle layer), and a compliance-matched TEVG (n = 6 aortas; n = 6-10 grafts). **c**. Pressure–outer diameter response (mean ± s.d.) for rat aorta and compliance-matched TEVG. **d**. Representative image of the compliance-matched TEVG implanted as rat abdominal aorta interpositional grafts. **e**. Standardized acquisition set-up (16x water-dip objective) and sample orientation (longitudinally) during acquisition in a 35 mm dish. **f**. Representative multiphoton images acquired at Ex: 792 nm showing imaging of the scaffold-only (top) and an 8wk double-joined artery-TEVG culture (bottom) in the three collection windows used throughout (395/25 nm for SHG, vs. 460/40 nm & 620/60 nm for 2PEF) with merged channels and 3D renderings. **g**. Voxel-intensity distributions of scaffold-only stacks (log-scaled voxel count) defining channel-specific background (dashed lines) in comparison to an 8wk culture sample.

### A compliance-matched TEVG as a label-free benchmark for longitudinal multiphoton imaging

To establish a label-free optical baseline for longitudinal artery-graft imaging, we first defined interpretable excitation and emission windows with rat aortic sections and freshly explanted aortas and confirmed that *de novo* SHG could be detected under non-destructive imaging conditions using diced aortic explants cultured for several weeks (**Extended Data Fig. 3a-d**). We then selected a trilayered compliance-matched TEVG as the reference graft for our *ex vivo* culture studies, enabling optical benchmarking against a companion rat abdominal aortic implantation study using the same graft design. This graft consisted of electrospun polyester urethane urea (PEUU) elastomer blended with porcine gelatin in the middle and outer layers, with a luminal sulfobetaine modified PEUU (PESBUU-50) layer for hemocompatibility (**Fig. 2a**)^33,34^. Iteratively, we adjusted layer thicknesses to tune passive circumferential compliance to match that of healthy *ex vivo* rat abdominal aorta across physiological pressure ranges (**Fig. 2b-d**).

**Fig. 3.**
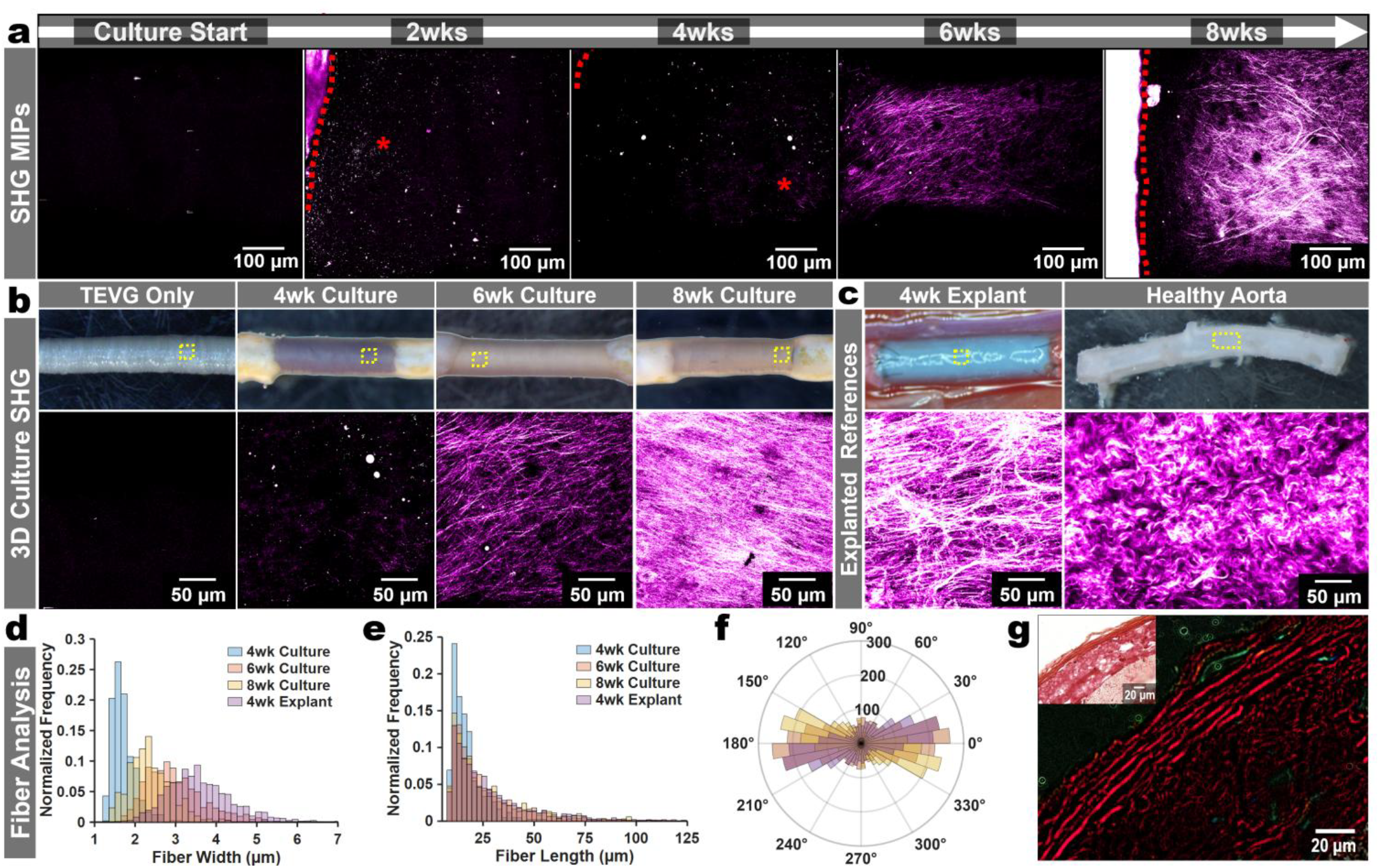
Multiphoton Longitudinal SHG Assessments. **a**. Representative SHG maximum-intensity projections acquired longitudinally from culture start through 8 weeks (792 nm excitation, 395/25 nm emission). Red dashed lines indicate where the aortic boundary is, and red asterisks (for 2wk & 4wk) indicate where SHG fibers are newly forming. **b**. Representative gross images (top) and corresponding higher magnification (bottom) of TEVG only and TEVGs cultured for 4, 6, and 8 weeks. **c**. Reference gross images (top) and corresponding SHG MIP images (bottom) of a 4-week (4wk) interpositional explant and a healthy aorta. Yellow dashed boxes indicate relative areas shown in SHG images below. **d**. CT-FIRE-based quantification of SHG-positive fibers shown as normalized frequency distributions of fiber width for the samples shown in b and the 4wk explant reference in c. **e**. CT-FIRE-based quantification of SHG-positive fibers shown as normalized frequency distributions of fiber length for the samples shown in b and the 4wk explant reference in c. **f**. CT-FIRE-based quantification of SHG-positive fibers shown as normalized frequency distributions of fiber angle for the samples shown in b and the 4wk explant reference in c. **g**. Picrosirius red staining of an 8wk cultured TEVG section with polarized-light imaging confirming collagen birefringence in collagen-rich regions.

We standardized an excitation and collection scheme based on the signatures from both tissue and our graft design and selected a 16x objective lens for all downstream work based on its signal transmittance and relative imaging depth penetration (**Extended Data Fig. 4a-c**). Using this, we found that acellular TEVGs exhibited modest intensity in the 460 nm channel, strong signal in the 620 nm channel, and minimal signal in the 395 nm (SHG) window (**Fig. 2e,f**). Single-channel views, merged images, and 3D reconstructions further supported the presence of fibrillar SHG and cell-associated 2PEF in cultured constructs that were absent in scaffold-only controls (**Fig. 2f**). Relative to scaffold-only controls, 8-week artery-TEVG cultures exhibited clear shifts in channel-intensity distributions across all three windows, indicating the emergence of endogenous remodeling-associated changes over scaffold background (**Fig. 2g**). As scaffold-associated 2PEF may vary due to polymer chemistry and various fabrication methods, we utilized several biomaterials (electrospun gelatin, PEUU, PESBUU-50, PCL, and UBM-hydrogel coated PCL) in our platform. After 4 weeks of culture, each material supported consistent longitudinal imaging and elicited specific differences in cell-associated 2PEF and SHG, with a limitation in materials with intrinsic SHG, such as urinary bladder ECM hydrogel coatings (**Extended Data Fig. 5**)^35^.

**Fig. 4.**
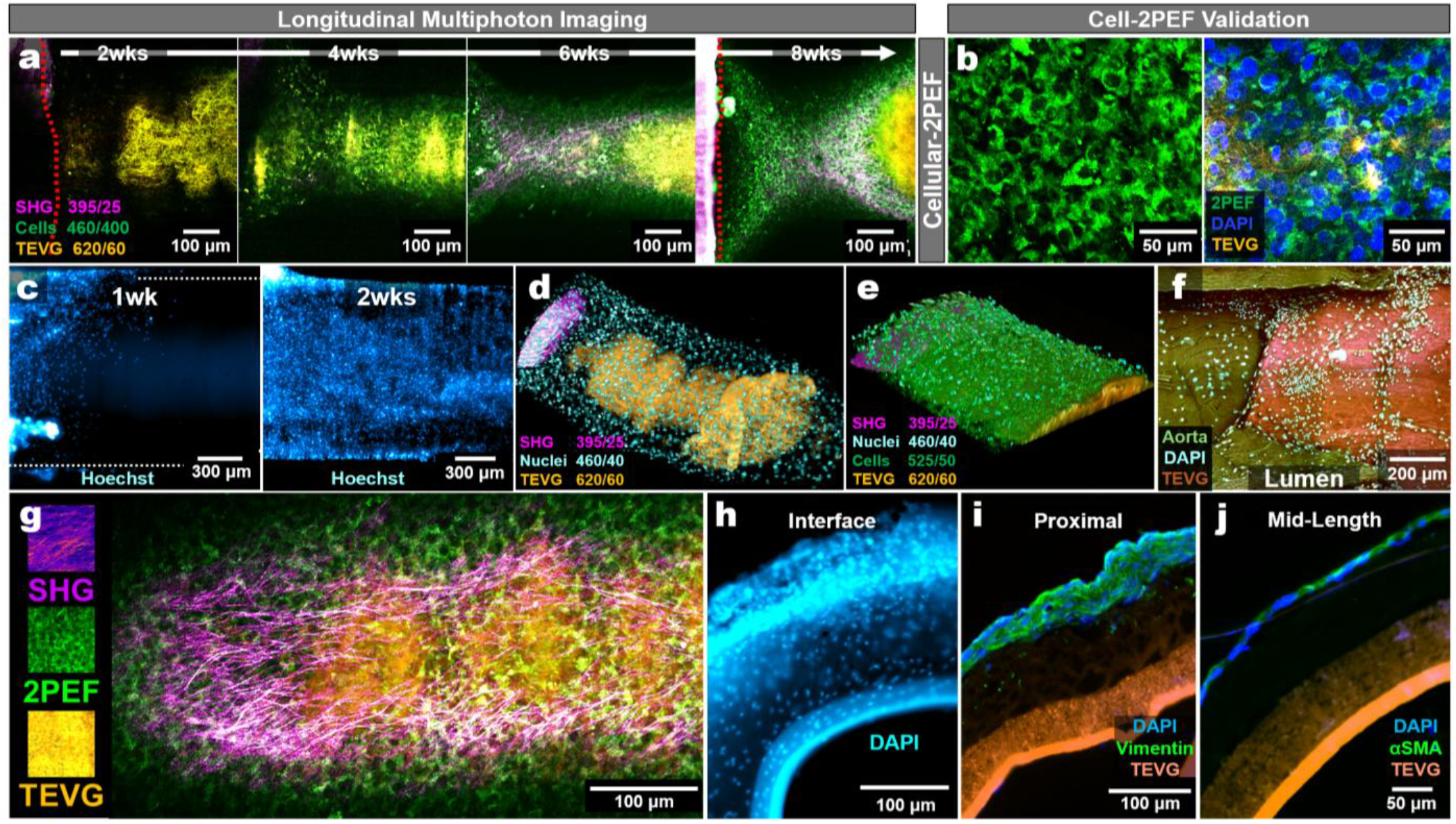
3D Culture Cellularization and Longitudinal Tracking. **a**. Longitudinal live two-photon imaging of TEVG cellularization in the 3D *ex vivo* culture over 2-8 weeks. Images are displayed as merged channels under 792 nm excitation, showing SHG (magenta), cellular two-photon excited fluorescence (2PEF; green), and TEVG background/autofluorescence (red) wherein overlapping red and green appear as yellow/orange. Red dashed lines indicate where the aorta is in view. **b**. (left) A representative multiphoton image from a 6wk culture sample showing endogenous 2PEF (Ex/Em: 792/460 nm) highlighting cell-like structures developing upon the TEVG surface. (Right) A 6wk culture that was imaged, DAPI stained, and then reimaged verifying cell-2PEF corresponds to nuclei. **c**. Widefield inverted fluorescence images of Hoechst-stained cultures at 1wk and 2wks to demonstrate potential for volumetric cell tracking. **d**. A 3D reconstructed view of Hoechst-stained nuclei across the TEVG after 2wks of culture. **e**. A 3D reconstructed view of Hoechst-stained nuclei across the TEVG after 6wks of culture. **f**. A fixed and DAPI stained sample was cut, and the inner lumen was imaged facing upward. DAPI staining required 720 nm excitation, shifting SHG off scale. Image shows the aorta (dark green, Em:460 nm), the TEVG (orange, Em:395 nm), and DAPI-stained nuclei (cyan, Em:525 nm). **g**. Representative high-resolution image at 16x showing an 8wk culture sample, able to distinguish cells, SHG, and the TEVG. **h**. Gross cross-section of a 2wk cultured TEVG (near the aortic interface) fixed and stained with DAPI to visualize nuclei. **i**. Immunostaining for vimentin. **j**. Immunostaining for α-SMA.

**Fig. 5.**
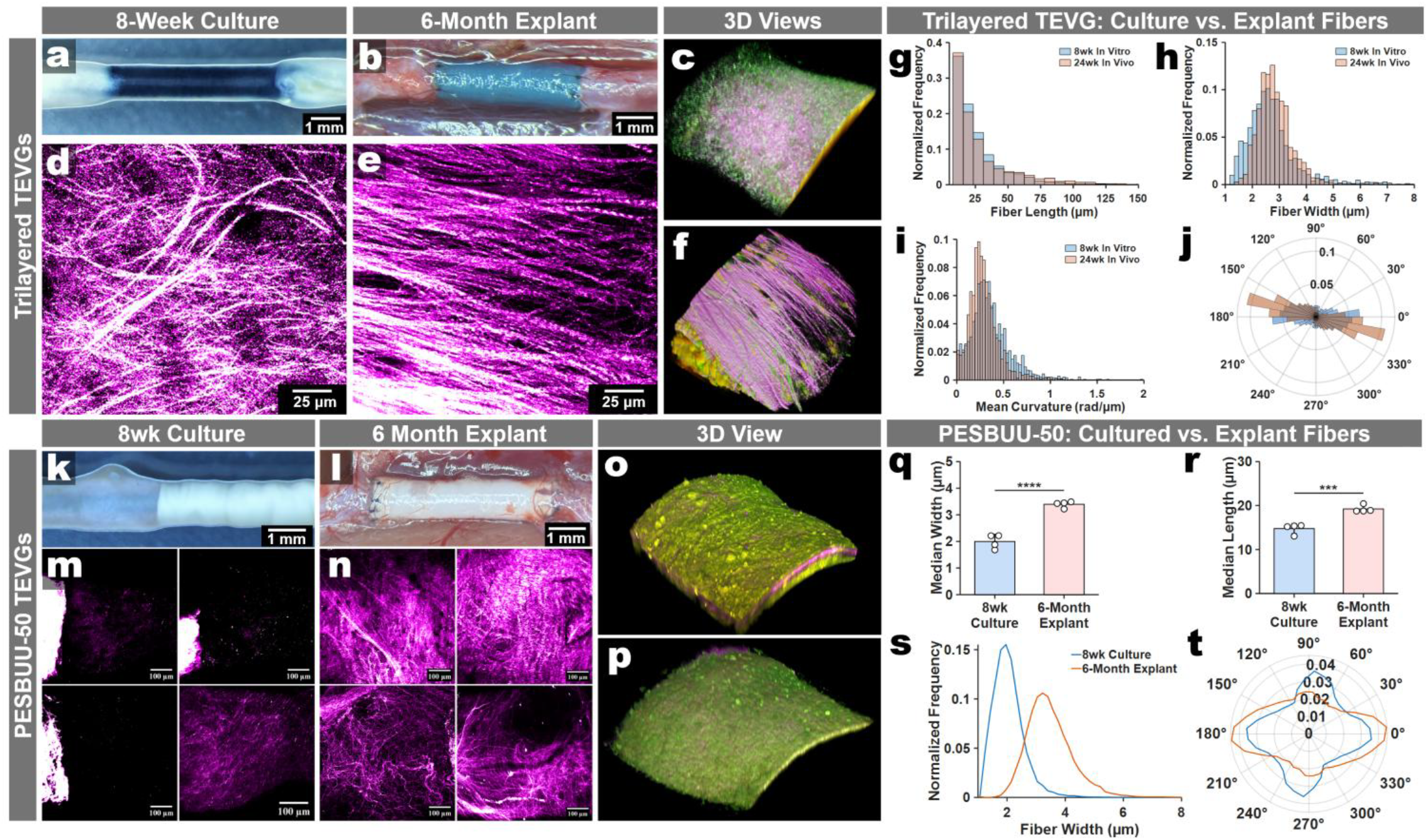
*Ex vivo* culture partially mirrors scaffold-dependent collagen remodeling associated with long-term implants. **Trilayered TEVGs: a**. Representative gross images of an 8wk cultured graft. **b**. The same graft design in situ at 6 months post-implantation immediately prior to explant. **c**. 3D reconstruction image of the 8wk culture. **d**,**e**. Second harmonic generation (SHG) maximum intensity projections (MIPs; magenta hot) from 8wk culture (**d**) and 6-month explant (**e**). **f**. 3D reconstruction image of the 6-month explant. **g-j**. Collagen fiber metrics quantified from SHG images (blue, 8 weeks *in vitro*; orange, 6 months *in vivo*): (**g**) fiber length, (**h**) fiber width, (**i**) mean straightness/curvature (rad/µm), and (**j**) fiber orientation. **PESBUU-50 TEVGs:** (**k**,**l**) Representative gross images of an 8wk cultured graft (**k)** and 6-month explant *in situ* (**l**). **m**,**n**. SHG MIPs of the 8wk culture (**m**) and 6-month (**n**) sample replicates. **o**,**p**. Representative 3D views of the 6-month explant (**o**) and 8wk culture (**p**). **q-t**. Corresponding SHG-derived fiber metric distributions (blue, 8wks *in vitro*; orange, 6-months *in vivo*): (**q**) fiber length, (**r**) fiber width, (**s**) mean overlaid fiber widths, & (**t**) mean fiber orientations. Statistics are one-way ANOVA with Tukey post-hoc, n = 4, ***P≤0.001 and ****P≤0.0001.

### Longitudinal multiphoton imaging resolves time-dependent collagen-associated SHG remodeling

After establishing that *de novo* fibrillar SHG could be detected on TEVG surfaces, we then investigated whether SHG could be monitored longitudinally throughout the culture duration. Across 8-weeks of culture, fibrillar SHG progressively increased along the graft surfaces, indicating a time-dependent accumulation of collagen structures (**Fig. 3a**). To quantify the collagen architecture, we applied the open-source SHG-fiber analysis pipeline CT-FIRE to characterize microstructural features such as fiber width, length, and straightness^36^. SHG-positive structures initially appeared in baseline artery-TEVG cultures by week 2 as short and thin microfibers, which became more distinct by week 4, and matured into longer and thicker fibrillar strands by weeks 6-8, resembling the architecture observed in 4-week interpositional graft explants (**Fig. 3b-f**).

Although signal intensity and resolution of the SHG was reduced with lower-magnification objectives, monitoring remained feasible, suggesting a potential route toward faster screening workflows (**Extended Data Fig. 6**). Because SHG collection is reported to be sensitive to sample orientation and geometry, we tested the effect of sample orientation upon the resulting SHG collection and subsequent fiber analyses and found a modest sensitivity to sample orientation during imaging, leading us to standardize the sample orientation across all studies (**Extended Data Fig. 7**)^37,38^. We further validated the presence of collagen where SHG signatures were apparent with picrosirius red staining of sectioned samples and confirmed birefringence of these areas under polarized light (**Fig. 3g**).

**Fig. 6.**
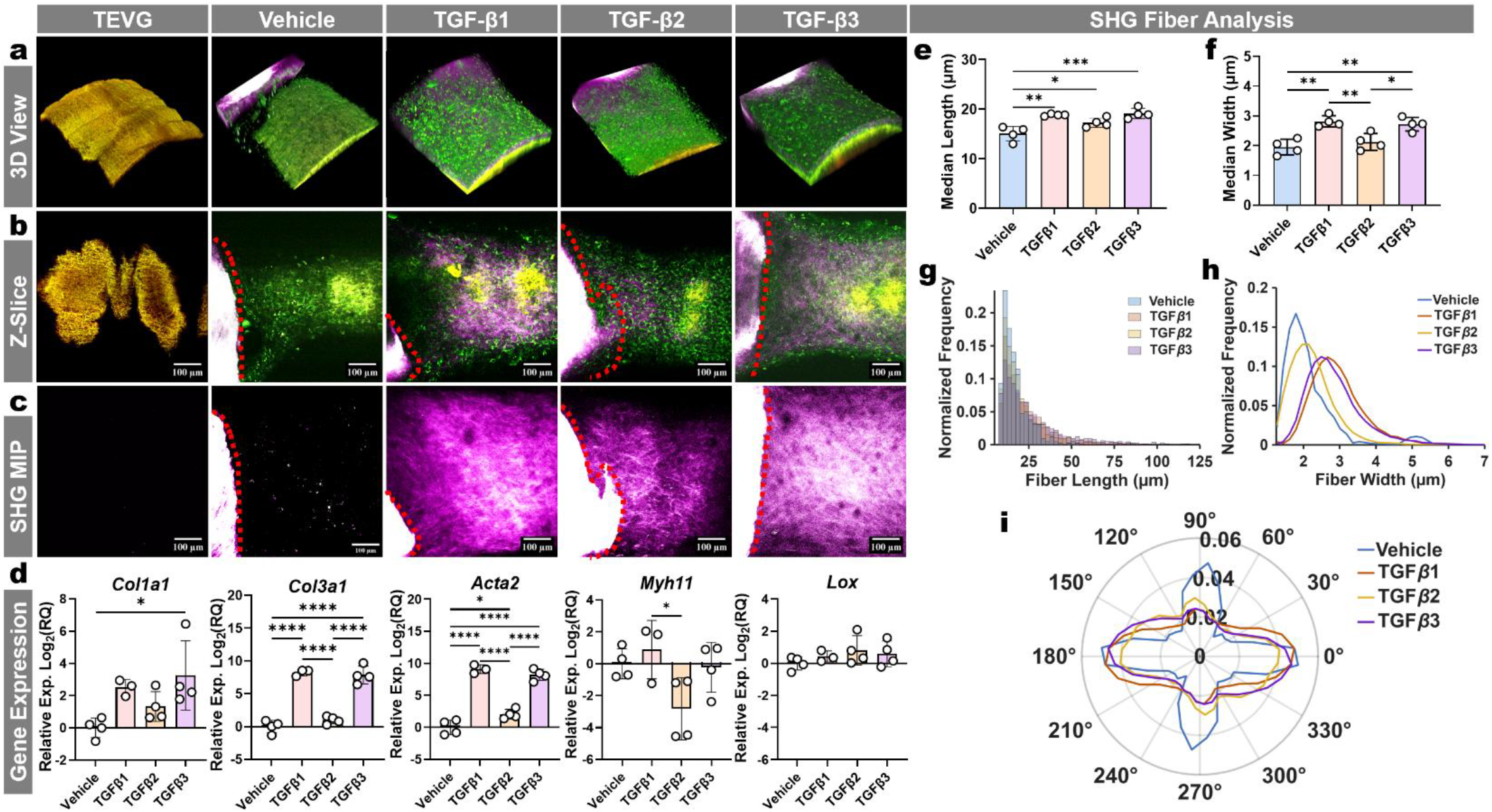
3D Culture Sensitivity to Exogenous TGF-β Isoform Treatments. **a**. Representative 3D reconstructions of 4wk cultured samples from each group. **b**. Representative z-slice images of 4wk samples stimulated with 10 ng/mL of rhTGF-β isoforms every three days. **c**. A representative SHG max intensity projection (MIP) from each group. Red dotted lines in (b) and (c) depict the aortic boundary. **d**. Gene expression levels of the rhTGF-β treated groups normalized to the vehicle treated group. Reference gene was *Rer1*, and results are depicted on a Log_2_(Fold-Change) scale. **e**. Collagen fibers were segmented from SHG (395/25 nm) channel MIPs using CT-FIRE, and fiber length and **f**. fiber width were quantified for each replicate. For each replicate, the median fiber length (**e**) and median fiber width (**f**) were calculated across all detected fibers, and these replicate medians were used for plotting and statistics. **g**,**h**. Plots of the normalized frequency of each group’s fiber lengths and widths (each replicate normalized to its respective total fibers). **i**. Polar plot shows the normalized distribution of collagen fiber orientation angles for each group. Fiber angles were binned (10° bins), replicate histograms were normalized to probability, and the group trace represents the mean across replicates. Statistics (qPCR) per gene are ordinary one-way ANOVA with Tukey post-hoc (n = 4). Statistics for (e) and (f) are ordinary One-way ANOVA followed by a Tukey post-hoc test (n = 4). *P≤0.05, **P≤0.01, ***P≤0.001, and ****P≤0.0001.

### Endogenous 2PEF enables longitudinal tracking of TEVG cellularization

Before fibrillar SHG became prominent, we observed progressive endogenous 2PEF signals in the 460/40 nm channel across TEVG surfaces, with morphologies consistent with cellular structures, suggesting that cellularization could be monitored in parallel with SHG-based remodeling (**Fig. 4a**). Alone, this 2PEF signature is commonly accepted as endogenous cellular 2PEF, however, our scaffold autofluorescence included signal in 460/40 nm. To establish that this signature was cell-associated, we imaged cultured samples, stained them with DAPI, re-imaged the same regions and overlaid registered ROIs. The 460 nm 2PEF signal surrounded the DAPI-positive nuclei, supporting its interpretation as a cell-associated signature in this platform (**Fig. 4b**). At early timepoints, however, cellular-2PEF was weak or absent despite the presence of cell nuclei on TEVG surfaces at 1 and 2 weeks by DAPI staining (**Extended Data Fig. 8**). Live Hoechst nuclear labeling confirmed that cells were present upon the TEVGs by 1- and 2-weeks, irrespective of cellular-2PEF intensity (**Fig. 4c**). As Hoechst signal diminished over time, we found that cellular-2PEF, SHG, TEVG-2PEF, and nuclear labeling could be captured together in parallel with remodeling (**Fig. 4d,e**).

Transection and imaging of DAPI-stained TEVG segments at the interface further confirmed nucleated cells along the lumen interface, with morphology consistent with host tissue extension onto the TEVG luminal surface (**Fig. 4f**). At terminal endpoints, we found that higher-resolution of these label-free readouts could be performed with fixed samples (**Fig. 4g**). We confirmed radial infiltration of cells into the TEVGs near the artery-graft interface and performed immunofluorescent staining on regions towards the center of the TEVGs, verifying the presence of vimentin and α-SMA expressing cells as expected from our culture media conditions (**Fig 4h-j**). The platform is also compatible with murine explants and genetically encoded fluorescent reporters. Using wild-type and Myh11-CreERT2;Rosa26-mTmG dual-reporter aortas, we resolved SHG, cellular-2PEF, and scaffold-2PEF as before and enhanced GFP emission (Ex: ∼900 nm, Em: 525/50 nm), supporting the integration of lineage-reporter imaging into the platform for broader utility (**Extended Data Fig. 9**).

### Remodeling trajectories in the culture model partially mirror long-term remodeling *in vivo*

We then probed whether the longitudinal collagen remodeling patterns observed in the culture platform corresponded to the general remodeling phenotypes and microstructural architecture found in long-term implantation two graft designs. For the compliance-matched trilayered TEVG, we compared SHG-defined fibrillar collagen architecture in 8-week cultures with that of 6-month rat abdominal aortic interposition explants acquired using our platform standardized imaging en face orientation (**Fig. 5a-f**). 3D reconstructed views prepared from imaged 3D stacks revealed that SHG was predominant in vivo and in vitro, appearing upon the outer surfaces of the grafts (**Fig. 5c vs. 5f**). Cultured grafts developed dense, aligned SHG-based collagen fibers that closely resembled the architecture observed in long-term explants. As before, we quantified collagen fiber organization using SHG-derived metrics (fiber length, width, straightness, and angular distributions) and found that the distributions between 6-month models and the 8-week cultures were highly similar in fiber length, width, straightness, and orientation, with the total amount of SHG signal being expectedly higher in explanted grafts (**Fig. 5g-j**).

We also compared the outcomes between implantation and culture for electrospun PESBUU-50 grafts. In contrast to the trilayered grafts, we found the highest levels of SHG-based fibers were locationally different. In both the 8-week cultured samples and in 6-month explants, we found a similar divergence between explants and cultures, but with similar trajectories in overall graft-specific trends (**Fig. 5k-n, Extended Data Fig. 10**). The trilayered grafts SHG each had SHG-signatures predominantly along the outermost surfaces, whereas both the PESBUU-50 explants and culture samples expressed their highest SHG-signatures beneath 2PEF-positive cells, closer to the graft surface (**Fig. 5o,p vs. 5c,f**). However, our quantitative analyses of the SHG-fiber showed significant shifts in median fiber width and length, with explanted samples retaining qualitatively and quantitatively more collagen deposition, again supporting that while trajectories align, in vivo implantations retain much higher levels of remodeling (**Fig. 5q-t**).

### The 3D culture model is sensitive to exogenous protein treatments

To demonstrate the system’s responsiveness, we evaluated whether SHG was impacted by human recombinant TGF-β isoform treatment, well-established inducers of collagen deposition in aortic culture contexts. Longitudinal tracking of each condition (n = 4 independent cultures per group) showed increases in SHG intensity at 4-weeks with TGF-β isoforms 1, 2, and 3 treatments (10 ng/mL every 3 days) relative to vehicle controls. Across 3D reconstructions, z-planes, and max intensity projections of SHG, we found a strong increase in SHG intensity and appearance with TGF-β isoform treatment **(Fig. 6a-c)**. To validate these results and verify that the culture system remains compatible with endpoint assays often used in vascular contexts, we performed qRT-PCR at 4 weeks and found that TGF-β isoforms altered expression of contractile and matrix-remodeling markers (including *Myh11, Acta2, Lox, Col1a1*, and *Col3a1*) relative to vehicle controls, supporting that the observed microstructural SHG differences are accompanied by significant differences in gene expression (**Fig. 6d**). We next quantified SHG fiber architecture using CT-FIRE. In comparison to vehicle treatment, TGF-β treatment shifted fiber-length and fiber-width distributions and altered group medians, demonstrating that the platform resolves remodeling at the level of collagen microstructure rather than relying solely on qualitative imaging (**Fig. 6e-h**). Notably, TGF-β2 produced narrower fibers than TGF-β1 and TGF-β3, consistent with its more modest induction of collagen-associated readouts (**Fig. 6f,h**). In contrast, fiber orientation distributions were broadly similar across treatment groups, suggesting that TGF-β primarily modulated fiber dimensions under these conditions rather than imposing a dominant alignment axis (**Fig. 6i**).

## Discussion

A major constraint in the development of small-diameter tissue-engineered vascular grafts is that functionally relevant remodeling outcomes are typically resolved only after surgical implantation, despite the time, cost and low iteration capacity of these studies. In this work, we developed a geometry-preserving ‘organotypic’ artery-graft culture platform that enables longitudinal, label-free multiphoton monitoring of host-graft remodeling over multiple weeks while maintaining a three-dimensional cylindrical interface resembling interpositional grafting. In contrast to endpoint-focused *in vivo* workflows and *ex vivo* systems that culture vessels or grafts separately, this platform permits repeated interface-focused imaging of cellularization and fibrillar matrix remodeling in an accessible, low-cost, intermediate design. By stabilizing artery and graft segments on matched holders without sutures, adhesives, or clamps, the platform avoids introducing additional anastomoses-related variables while preserving the option to incorporate them systematically in future studies. The platform’s sensitivity to dimensional matching further underscores the importance of local geometry at the anastomosis, consistent with evidence that mismatch can influence patency and participate in maladaptive wall remodeling associated with intimal hyperplasia^39^.

A central advance of this work is that the platform supports longitudinal visualization and additionally generates time-resolved microstructural readouts that potentially forecast the trajectories in long-term graft implantation remodeling outcomes. While some organotypic vascular culture models exist, these are largely limited to terminal endpoints and macroscale ultrasound imaging for longitudinal readouts^18-25^. We expand upon these by utilizing label-free multiphoton imaging to track the emergence of endogenous cell-associated 2PEF, *de novo* fibrillar SHG-based collagen on graft surfaces across culture duration, allowing us to quantify evolving collagen architecture using established and publicly accessible SHG-fiber analysis tools. Our orthogonal histology supports the interpretation of SHG-positive structures as collagen-rich remodeling features, and our nuclear co-registration and live cell labeling support the interpretation of this 2PEF signal as a cell-associated readout. Importantly, drawing from parallel studies using the same compliance-matched trilayered TEVG design for animal implantation, we found that the collagen architecture observed after 8 weeks in culture was in strong agreement with those found in 6-month interpositional explants. This concordance across fiber length, width, straightness and orientation supports the concept that the culture platform captures aspects of remodeling trajectories that are relevant to long-term *in vivo* microstructure, whereas monolayer cultures are often limited to disordered matrix deposition. The PESBUU-50 graft provided a case wherein both the *ex vivo* culture and long-term explants exhibited minimal SHG deposition overall, signifying a generalized match between the trajectory of the culture and the long-term remodeling states. Investigating samples with the highest SHG levels, we continued to find partial fiber correspondence, signifying a second graft design with similar outcomes in the platform with those from long-term implantation. Together, these findings support the platform as a decision-informing intermediate model that mirrors many aspects of long-term graft remodeling trajectories, and broader validation across a range of future studies will ultimately define its full predictive scope.

Using the platform, we resolved biologically induced differences in remodeling from TGF-β isoform treatments. While prior studies have shown that the three TGF-β isoforms can drive distinct remodeling in different contexts using knock-out models, vascular remodeling studies comparing differences after treatment with each isoform remains limited. We found that treatment with TGF-β isoforms increased SHG-positive remodeling relative to vehicle conditions and produced accompanying shifts in the expression of contractile and matrix-remodeling genes, supporting that the structural changes observed by imaging reflected altered biological states. Importantly, SHG-fiber analyses found that these perturbations were not equivalent, as isoform treatment led to distinct and significant microstructural distributions, paralleling the relative gene expression between the isoform treatment groups. This perturbation responsiveness potentially positions the platform as a controllable experimental system in which biochemical cues can be linked to evolving matrix architecture and downstream molecular endpoints.

This platform has several limitations that define its intended use. Our model lacks systemic contributors from blood circulation, such as various infiltrating immune and progenitor populations, innervation, and multi-organ signaling. Similarly, because the platform is static, it does not incorporate established drivers of vascular remodeling such as hemodynamic flow or axial loading, nor does it monitor or maintain endothelial function. This is an intentional trade-off from prioritizing configurations conceptually closer to established models such as the aortic ring assay, specifically to promote accessibility and higher-throughput parallelization. Additional limitations include the use of SHG as a robust indicator of fibrillar collagen organization as prior reports have verified that SHG is sensitive to sample orientation, cannot directly quantify total collagen content, and is likely to underrepresent very thin, disordered or non-fibrillar matrix features^40,41^. Likewise, optical signatures intrinsic to a scaffold can constrain monitoring of collagen remodeling for some biomaterials, as illustrated by residual SHG in UBM-coated PCL constructs. For applications requiring definitive total collagen quantification or compositional profiling, SHG quantification should be complemented by established orthogonal biochemical assays. Lastly, the platform’s use of multiphoton imaging may arguably limit accessibility for some investigators. Although optimized for multiphoton microscopy, those lacking access to adequate instrumentation can potentially adapt the platform for widefield imaging with live-cell dyes or murine fluorescent reporter strains, and longitudinal collagen monitoring can be replaced with serial endpoint assays. Further, although we did not evaluate terminal whole-mount clearing, staining, and light-sheet microscopy here, recent work suggests that these approaches could complement the platform by providing high-resolution volumetric insight into matrix remodeling, cellular distribution, and three-dimensional tissue organization across the entirety of the samples^42^.

Taken together, this organotypic artery-graft culture platform, coupled with longitudinal multiphoton tracking, establishes a practical bridge between overly simplified monolayer *in vitro* screening and resource-intensive implantation studies. The platform’s compatibility with multiple scaffold classes, rodent (mouse and rat) explants, and genetically encoded reporters suggests that the system can be adapted to broader mechanistic questions, including cell lineage contributions to interface remodeling and the effects of biomaterial properties upon early graft integration. Our findings support the platform as an intermediate model for pruning hypotheses, probing mechanistic pathways, and comparing graft remodeling trajectories before and in parallel with animal implantation. More broadly, the platform enables direct perturbation and longitudinal tracking of microstructural remodeling over time at geometrically relevant artery-graft interfaces, allowing investigation of processes that remain difficult to resolve in conventional implantation studies and prior culture models.

## Methods

### Detailed protocols and analysis code

The detailed protocols for the 3D culture model, multiphoton imaging, and all MATLAB code for plotting are available online.

### Materials and custom components

Reagents, consumables, equipment, and vendor catalog numbers are provided in **Extended Data Table 1**. Hexagon-shaped 3D-printed polycarbonate (PC) sample holders were used to provide consistent height alignment and stable tissue-TEVG contact and allow rotation of the samples, as needed. The hexagonal designs consisted of a 5 mm x 5 mm x 5 mm shape with a 0.5 mm hole at the center. 3D printing was performed using an Ultimaker Pro 5. Printing settings were set to an infill of 100% using transparent PC (Ultimaker, USA). This design is available as a downloadable .STL file.

After printing, PC holders were inspected and cleaned with sterile DI water, soaked in 70% EtOH for 4 h to remove any free unbound polymers, and then sterilized by autoclave at 121°C while on a 500 μm OD mandrel. Before use, the holders were placed in sterile 1x DPBS overnight and rinsed in sterile 1x DPBS immediately prior to use. PC holders are disposable, but reuse is feasible with sterilization. PTFE-coated stainless-steel wire mandrels with a 500 µm outer diameter (OD) were cut to 30 mm lengths and used to support and align the samples above the culture surface. Mandrels were sterilized by autoclaving at 121 °C. Mandrels were intentionally extended beyond the sample holders to enable handling with sterile forceps during assembly and transfer.

### Culture medium preparation

The presented culture system utilizes complete Smooth Muscle Cell Growth Medium (SmGM-2; Lonza, Germany). Complete media was prepared according to the manufacturer’s instructions and supplemented with the proprietary SingleQuots SmGM-2 vials which include 25 mL of FBS (5%), insulin, hFGF-B, hEGF, and GA-1000 (gentamicin sulfate-amphotericin). At the time of this publication, the vendor verified that SmGM-2 is based on MCDB 131 media containing 5.55 mM glucose, 1 mM pyruvate (pyruvic acid), and 10 mM glutamine. As MCDB 131 media was originally formulated to include trace elements, we confirmed with the vendor that SmGM-2 includes 0.005 µM cupric sulfate pentahydrate (1.25 µg/L). Complete media was aliquoted into sterile vials and stored at 4°C in a light-protected box and was used within 1 month of preparation. Before changing, aliquoted media was covered with a gas-permeable sterile membrane film and placed into a CO_2_ incubator (37°C, 5% CO_2_) for 30 mins prior to use to stabilize pH and temperature.

### Rat and mouse aortic explant isolation

All studies were performed under IACUC approved animal care protocols. Lower abdominal aortas were harvested from young adult male Sprague Dawley rats (6-8 weeks old, ∼200 g). Prior to explant, animals were sprayed with 70% EtOH before entry into the biosafety cabinet. Using pre-sterilized tools, the abdomen was opened to expose the aorta, and organs were gently displaced to minimize rupture and bleeding. The aorta was clamped above the aortic bifurcation and below the infrarenal region to obtain ≥ 2 cm of vessel. To minimize exposure to air, the aorta and vena cava were explanted together and submerged in 4°C sterile 1x DPBS supplemented with 2% penicillin-streptomycin and 2% amphotericin B. These were separated under a stereomicroscope and gently flushed to remove any residual blood and clots. Aortas were handled consistently and cleared of peri-adipose tissue using sterile forceps, with care to minimize damage to the adventitia of the vessel. The total time from euthanasia to final placement in the incubator was maintained at ≤ 45mins per sample (average time ∼30 mins). Longer processing times (>1 hr) led to reduced cell growth.

For murine wild-type explants, adult male C57BL/6J mice were purchased from Jackson Laboratories. For smooth muscle lineage tracing, B6.FVB-Tg(Myh11-icre/ERT2)1Soff/J mice (JAX Stock# 019079) were crossed with B6.129(Cg)-Gt(ROSA)26Sor^tm4(ACTB-tdTomato,-EGFP)Luo/J reporter mice (Rosa26-mTmG; JAX Stock# 007676) to generate Myh11-CreERT2;Rosa26-mTmG dual reporter mice. To induce CreER^T2^-mediated recombination in MYH11-expressing smooth muscle cells, tamoxifen was administered at 75 mg per kg of animal mass at a concentration of 20 mg/mL for 5 consecutive days via intraperitoneal injection in filtered corn oil (Millipore Sigma). Following induction, Cre-mediated recombination switches reporter expression from membrane tdTomato to membrane GFP in recombined cells. Mice were harvested within 5 days after the final tamoxifen dose.

### TEVG fabrication and preparation

In the presented method, we use a trilayered electrospun graft composed of elastomers that have been previously published including polyester urethane urea (PEUU) and its 50% sulfobetaine carrying variation, PESBUU-50^33,34,43,44^. In both cases, these polymers were prepared in-house as described in Ye, et al. from 2014^33^. This trilayered design consists of an inner PESBUU-50 layer, a middle PEUU:Gelatin (80:20) layer, and an outer PEUU:Gelatin (20:80) layer. This specific TEVG design was chosen to parallel our ongoing rat aortic interpositional implantation studies, and PESBUU-50’s specific role was to provide an antithrombogenic elastomeric inner layer. The TEVGs were electrospun using a temperature, voltage, and humidity-controlled IME electrospinning system (Vivolta, Netherlands). The settings to fabricate the TEVGs are included in **Extended Data Table 2**. The trilayered TEVGs were crosslinked in 0.5% genipin in 200 proof EtOH for 24 h at 37°C and washed 3x in 200 proof EtOH. Prior to use, TEVGs were sterilized with 70% EtOH, rinsed 3x in sterile 1x DPBS, incubated in sterile 1x DPBS for 12 h. Samples were then rinsed in sterile 1x DPBS immediately before use.

### *Ex vivo* circumferential mechanical compliance testing

Mechanical compliance testing was performed on a closed-end tubular biaxial mechanical loading device (CellScale Biomaterials Testing, Waterloo, Canada). Characterization of rat abdominal aortas was performed *ex vivo* with testing parameters matched between explanted aortas and electrospun scaffolds. Prior to testing, aortas were explanted, cleared of peri-adipose tissue, and flushed clear of blood clots. Aortas and TEVGs were washed with 1xDPBS heated to 37°C and allowed to acclimate for 20 mins prior to testing. Prior to testing, the tubular mechanical testing device was set-up by focusing the camera to accentuate the outer diameter of the samples and assigning size/scale calibration. The sample bath was filled & maintained with 37°C 1x DPBS (aortas in 1x Krebs-Henseleit passive buffer to deactivate vSMCS) and acclimated for 30 mins. The system was then loaded with empty sample holders & load/pressure were tared. The aorta or TEVG was cannulated to sample holders using suture thread upon the ends of the holders and fully sealed from leakage using UV curable glue. Each sample was then loaded & tested using the parameters in **Extended Table 3**. Preconditioning was set for 9 cycles, and the 10th cycle was recorded as the test.

Compliance Calculations: Data was collected across taut axial stretch to match the axial loads in surgical implantation (generally, λ = 1.3-1.5 for aortas, and further defined by an axial load of 0.10 ±0.02 N. Compliance for our studies was defined as the mean ± 1σ of tested rat aortas at identical conditions. Compliance is calculated for each λ per bio-replicate, computed over ranges of 0.1 mmHg through 120 mmHg by the equation displayed at the bottom of **Extended Data Table 3**. [(OD120-OD70)/OD70]/(120mmHg – 70mmHg), which yields the ratio of the outer diameter (OD) differences when the lumen is pressurized at 70- vs. 120 mmHg. Results are reported as means ± 1σ. Tests with temp. drift >2°C, pressure >2 mmHg & slip/buckling were excluded.

### 3D artery-TEVG culture assembly and maintenance

The complete 3D culture constructs were assembled by combining segments of aorta (each 5 mm length; OD ∼1.2 mm, and wall thickness ∼0.10 mm) positioned adjacent to a 5 mm-long TEVG with OD/ID closely matched to the aortic segments, both upon a 500 µm polytetrafluoroethylene (PTFE)-coated mandrel. The wire mandrel with construct is then suspended on height-matched 3D printed PC holders placed at each end of the mandrel to maintain tissue alignment and consistent tissue-TEVG contact. Holder spacing could be adjusted to maintain contact if mild axial tissue contraction occurred during early culture. 100 mm and 60 mm dishes were used for isolation/cleaning steps, following transfer to 6-well plates in 4 mL of complete media for culture. Samples were maintained in standard culture conditions (37 °C, 5% CO_2_).

The lowest point of the assembled culture sample was no more than ∼3 mm below the air-media interface. Media was changed every 3 days, and samples were visually inspected daily during the first week to ensure tissue-TEVG contact. If contraction reduced contact between the aorta and TEVG within the first week, the holders were slid inward using sterile tweezers. Unless otherwise indicated, constructs were not joined using sutures, adhesives, or clamps to isolate remodeling outcomes attributable to tissue-TEVG apposition as these connection strategies may be introduced as independent experimental variables.

### Live imaging handling and aseptic practices

For live imaging, samples were handled in a biosafety cabinet to remove them from culture media, briefly washed in pre-heated 37 °C sterile 1x DPBS, and transferred to a 35 mm dish of 37 °C FluoroBrite DMEM supplemented with 1% Pen/Strep and 1% gentamicin. For optimal sample imaging conditions, we used an on-stage incubator system (Oko-Labs, Netherlands). We also verified that a culture incubator is not required and that reagents, such as Live Cell Imaging Solution (Thermo Fisher, USA) can be used for stable room-temperature imaging without CO_2_. Post-imaging, samples were immediately covered and returned to a biosafety cabinet, rinsed in pre-heated 37 °C sterile 1x DPBS, and placed back into standard culture conditions. Samples were imaged at 1-week intervals for ≤ 45 mins per sample. Across studies, we found that the most common risk of culture contamination was from the water-dip objective lens. This can be deterred by vigilant disinfection of the objective lens and the surrounding stage with the appropriate cleaners before and after every use.

### Multiphoton microscopy system hardware and device preparation

Multiphoton microscopy was performed using a TriM Scope II (Miltenyi BioTec, Germany) coupled to an Olympus BX51 upright microscope and a Ti:sapphire femtosecond InSight DeepSee+ Dual laser (Newport, USA). Acquisition and hardware control were performed using manufacturer provided software (ImSpector Pro, v7.5.0). Emission was collected with non-descanned PMTs in a backward (epi) configuration, and images were saved as OME.TIF files. We provide **Extended Data Figure 4** for more details on system configuration. Prior to imaging, the system was allowed to equilibrate for at least 20 mins. Desired laser power output was then measured for 5 mins using identical settings and plane as the samples. To minimize the risk of contamination from imaging sessions the microscope stage surfaces and the media-contacting portions of the objective were cleaned at least three times with 70% EtOH. The objective lens was cleaned with 200-proof biological grade EtOH and wiped with lens cleaning tissue (Whatman, Grade 105).

### Second harmonic generation (SHG) and two-photon excited fluorescence (2PEF) imaging

For histological slides, adventitial collagen in rat aortic sections was confirmed to generate SHG using 770-810 nm excitation with 377-395 nm emission detection, consistent with prior reports^45-47^. For live 3D cultures, SHG was acquired at 792 nm excitation and collected using a 395/25 nm emission filter. Using the same excitation wavelength, endogenous 2PEF was collected in parallel through 460/40 nm and 620/60 nm channels; an optional 525/50 nm channel yielded qualitatively similar cell-associated signal to 460/40. In merged images, TEVG autofluorescence was identified by signal in both 460/525 and 620 channels, whereas cells were predominantly enriched in the 460/525 channel^29,30,45^. In this work, 2PEF intensity was used as a cell-associated autofluorescence signal to support visualization/segmentation and was not interpreted as a definitive metabolic readout.

For SHG thresholding, native aortic tissue served as a positive control and TEVG-only cultures served as negative controls to define signal-to-noise-based thresholds. Live imaging settings were selected based on prior reports and practical experience for low-photodamage longitudinal imaging (≤15 mW at the sample, ≤5 µs pixel dwell time, and ≤30 s acquisition per z-plane)^26-30,48-50^. We provide **Extended Data Table 5** for our live imaging settings.

### Widefield fluorescence and birefringence imaging

Inverted fluorescence imaging was performed on an EVOS M7000 (Thermo Fisher Scientific, USA) using the DAPI v2, GFP v2, and RFP v2 filter cubes. Upright fluorescence and picrosirius red birefringence imaging were performed on a Nikon Eclipse 90i using an in-house linear polarizer. Further details and imaging settings are provided in **Extended Data Table 6**.

### RNA isolation and qRT-PCR

For gene expression assays, the TEVG portion of the constructs (excluding the original aortic tissue) were rinsed in sterile 1x DPBS and stored in RNAlater (Ambion) at 4°C for ≥24 h, transferred to TRIzol (Invitrogen), and stored at -80°C. Samples were homogenized on ice using Biomasher II tubes with a motorized pestle. Total RNA was isolated by TRIzol/chloroform extraction, and the aqueous phase was purified using the RNeasy Micro Kit PLUS (Qiagen). RNA concentrations were measured with a NanoDrop One (Thermo Fisher Scientific). RNA inputs were normalized and reverse transcribed using SuperScript IV VILO Master Mix (Invitrogen). Quantitative RT-PCR was performed on a QuantStudio 3 (Applied Biosystems, USA) using TaqMan single-plex MGB-FAM and TaqMan Gene Expression Master Mix in 20 µL reactions (MicroAmp Optical 96-well plates) using manufacturer-recommended cycling conditions. Relative-fold expression was calculated using the Livak ΔΔCt method (2^-ΔΔCt^), normalized to controls using the most consistent endogenous reference gene found (*Rer1*), and reported as relative quantification (RQ)^51^.

### SHG fiber quantification and image processing

All initial image processing was performed using ImageJ (FIJI) using version 1.54r, with some downstream assessments in MATLAB. As image post-processing techniques can vary across investigators, we utilized a previously published SHG fiber analysis application known as CT-FIRE (v3.0) to assess maximum intensity projections (MIP) taken from the (Ex: 792 nm) 395/25 channel z-stacks. CT-FIRE was used to extract SHG-positive fibers and quantify fiber width, length, orientation (0-180°), and straightness. CT-FIRE outputs were processed with custom MATLAB scripts to remove invalid entries, convert values to physical units using the imaging pixel size, apply consistent binning/ranges across groups, and compute per-sample summary values with probability-normalized histograms (bin counts divided by the total number of fibers per sample) to enable distribution-shape comparison across groups. All other image post-processing was performed using ImageJ/FIJI. CT-FIRE-based collagen fiber quantification (SHG). For all CT-FIRE analyses, we modified the following two settings from default: minimum fiber length: 15 pixels and maximum fiber width at 30 pixels.

In **Figure 6**, each sample (biological replicate) was individually analyzed, and CT-FIRE outputs were exported as histograms/arrays describing fiber length, width, angle, and straightness. For downstream analysis, CT-FIRE histogram CSV files (HistLEN, HistWID, HistANG, HistSTR) were batch-processed using a custom MATLAB script to harmonize binning across replicates and to generate replicate-level summary metrics. CT-FIRE outputs (pixel units) were converted to microns using the imaging calibration factor (µm/pixel). All statistical analyses were performed on replicate-level summaries (one value per sample) rather than pooled fibers. For each replicate, the median fiber length and median fiber width were calculated across all fibers detected within that sample. Fiber straightness was summarized as the replicate median of the straightness distribution (end-to-end distance/contour length and bounded between 0 and 1). Fiber alignment was quantified as a nematic order parameter (S), computed from the fiber angle distribution as (θ in radians; nematic periodicity 0-180°), yielding values from 0 (isotropic) to 1 (perfect alignment). The number of fibers contributing to each replicate metric was recorded for quality control. Angle distribution visualization (polar plots). To visualize orientation distributions across groups, fiber angles were binned in 10° increments and normalized per replicate to a probability distribution (each replicate contributed equally). Replicate distributions were averaged within each experimental group to obtain a group mean distribution. As collagen fiber orientation is nematic (0-180°), the one-sided distributions were mirrored to 0-360° for polar visualization graphs.

Prior reports have noted that collagen-based SHG can be polarization and orientation dependent, such that rotating a sample with anisotropic fibers by 90° relative to a fixed excitation polarization can alter SHG intensity and shift the apparent fiber-angle distribution by ∼90° ^37,38^. We verified subtle differences in fiber quantification and SHG intensity when samples were imaged at a ∼90° orientation with the same acquisition settings **(Extended Data Fig. 7)**. While these were relatively small changes with the samples remaining globally comparable, it motivated us to assign a single sample orientation for all images. We standardized our sample imaging orientation to be at 0° (placed with the length of the sample in a left-to-right orientation with respect to the microscope), and we recommend users of this methodology to standardize identifying their system’s polarization axis and assign a single specimen orientation across all imaging sessions included in a single study.

### Transforming Growth Factor Beta (TGF-β) isoform studies

For the TGF-β studies, carrier-free human recombinant (rh)TGF-β isoforms were reconstituted following the manufacturer’s recommendations at 20 µg/mL in sterile 4 mM HCl, aliquoted, stored in -20°C, with minimal freeze-thaw cycles per aliquot. All isoforms were purchased with matched vendor reported bioactivity levels. Treatments were provided at a concentration of 10 ng/mL of TGF-β into 4mL of media and vehicle controls were given an equal volume (2 µL) in the equivalent 4 mL of media.

### Preparation of alternative TEVGs used in the 3D culture

Polycaprolactone (PCL), PEUU, PESBUU-50, and gelatin TEVGs were electrospun in-house using the previously mentioned system. Electrospinning settings are provided in **Extended Data Table 7**. To deter degradation, gelatin TEVGs were crosslinked using 0.5% genipin for 24 h at 37°C. For urinary bladder matrix (UBM) hydrogel grafts, an inner 100% PCL sheet was first prepared by spin-casting a 10% w/v PCL (80,000 Mn) in HFIP solution onto glass to a thickness of ∼25 μm. UBM was prepared by the Hussey laboratory as previously described^52^. Porcine urinary bladders from market-weight animals were acquired from Tissue Source, LLC. Briefly, the tunica serosa, tunica muscularis externa, tunica submucosa, and tunica muscularis mucosa were mechanically removed. The luminal urothelial cells of the tunica mucosa were dissociated from the basement membrane by washing with deionized water. The remaining tissue consisted of basement membrane and subjacent lamina propria of the tunica mucosa and was decellularized by agitation in 0.1% peracetic acid with 4% EtOH for 2 h at 300 rpm on an orbital shaker. The tissue was then extensively rinsed with PBS and sterile water. The UBM was then lyophilized and milled into particulate using a Wiley Mill with a #60 mesh screen. To prepare UBM-hydrogels, we followed previously published methods^53^. Briefly, decellularized porcine UBM powder was solubilized at a concentration of 20 mg/mL in a solution of 1.0 mg/mL pepsin in 0.01 N HCl, with constant stirring for 24 h. The digest was then neutralized with 0.1 N NaOH, 1x PBS, and 10x PBS to a pH of ∼7.4, and a final concentration of 10 mg/mL. The samples were incubated at 37°C for 1 h for gelation, and then immediately applied to the PCL sheets as a thin uniform layer using a cell scraper in a biosafety cabinet. The UBM hydrogel-coated sheet was then manually rolled three times around a 1.1 mm diameter mandrel with coated surface facing outward. The seam was quickly sealed by pressing one side onto a 60 °C hot plate for <1 s.

### Statistical and Reproducibility

Statistical analyses and plots were performed in MATLAB. For gene expression results: For each gene, Log_2_(RQ) values were compared among groups using ordinary one-way ANOVA. Normality was assessed on the Log_2_ scale by Shapiro-Wilk test and homogeneity of variance by Brown-Forsythe test. Multiple comparisons used Tukey’s post-hoc test. Two-sided α = 0.05. Data are presented as mean ± SD with individual biological replicates shown. Technical replicates were averaged prior to analysis. For CT-FIRE fiber analyses, replicate medians (one point per biological replicate) were plotted by group in GraphPad Prism. Group summaries were displayed as mean ± SD (as specified) across replicate medians, with individual replicate points overlaid. Group differences were tested using one-way ANOVA with Tukey’s multiple comparisons, with statistical significance defined as p < 0.05.

## Acknowledgements

This research was funded by the National Institutes of Health (NIH) award 5R01HL157017 to JPVG. Additional funding was provided for DRM by a Supplement Award (S1) to the parent NIH R01HL157017. Additional support for MA was provided by P30GM131959, R41HL172481 and R21ES037105. Additional training support was provided for KMM by NHLBI F31 HL176056 and NHLBI T32 HL076124 (Cardiovascular Bioengineering Training Program). Graphical images were made using BioRender using the University of Pittsburgh’s institutional license. The authors would like to thank Pang-Wei Liu, Colleen O’Malley, and Dr. Keishi Kohyama for their assistance during the early phases of this project.

## Author Contributions

Conceptualization: JPVG, DRM; Methodology: JPVG, DRM, MA, and WW; Investigation: DRM, T. Murphy, T. Moyston, LXM, KMM; Data Curation: DRM; Formal analyses: DRM, JPVG; Software: JPVG; Resources: JPVG, G.S.H., WRW; Supervision: JPVG; Funding acquisition: JPVG; Writing - original draft: DRM; Writing - review & editing: all authors.

