## Extended Data Figures and Tables for "Resolving differential vascular graft remodeling using longitudinal multiphoton tracking in a 3D culture platform"

[1]Department of Bioengineering, University of Pittsburgh, Pittsburgh, PA, USA
[2]Department of Medicine, University of Pittsburgh, Pittsburgh, PA, USA
[3]Department of Surgery, University of Pittsburgh, Pittsburgh, PA, USA
[4]McGowan Institute of Regenerative Medicine, University of Pittsburgh, Pittsburgh, PA, USA
[5]Department of Cell Biology & Anatomy, School of Medicine, University of South Carolina, Columbia, South Carolina, USA
[6]Department of Biomedical Engineering Program, University of South Carolina, Columbia, South Carolina, USA
[7]Department of Chemical and Petroleum Engineering, University of Pittsburgh, Pittsburgh, PA, USA
[8]Department of Mechanical Engineering and Materials Science, University of Pittsburgh, Pittsburgh, PA, USA
[9]Vascular Medicine Institute, University of Pittsburgh, Pittsburgh, PA, USA
✉

#### Extended Data

1. **Extended Data Figures**
Extended Data Figure 1: An *Ex Vivo* Aortic Injury and Autologous Graft Reference
Extended Data Figure 2: 3D Culture Aorta-Graft Dimensional Matching
Extended Data Figure 3: Multiphoton Imaging of Aortic Sections & Diced Cultures
Extended Data Figure 4: Multiphoton Optical Configuration and Objective Lens Depths
Extended Data Figure 5: 3D Culture Platform Compatibility with Common TEVG Biomaterials
Extended Data Figure 6: Tracking SHG with a 4x Objective Lens
Extended Data Figure 7: Sample Orientation Impacts SHG Outcomes
Extended Data Figure 8: Fixed Sample 2PEF Validation with DAPI Staining
Extended Data Figure 9: Murine & Reporter-Strain Compatibility in the 3D Culture Platform
Extended Data Figure 10: PESBUU-50 6-Month Explants vs 8-Week Cultures.

2. **Extended Data Tables**
Extended Data Table 1: Reagents and supplies
Extended Data Table 2: Electrospinning parameters for trilayered TEVGs
Extended Data Table 3: Tubular mechanical compliance testing
Extended Data Table 4: Optical filters, PMTs, and imaging settings
Extended Data Table 5: Live imaging settings (16x Objective)
Extended Data Table 6: EVOS M7000 inverted widefield microscope configuration
Extended Data Table 7: Alternative biomaterial TEVG fabrication settings

3. **.STL File for 3D Printable Hexagon Sample Holders**

#### Extended Data Figure 1:

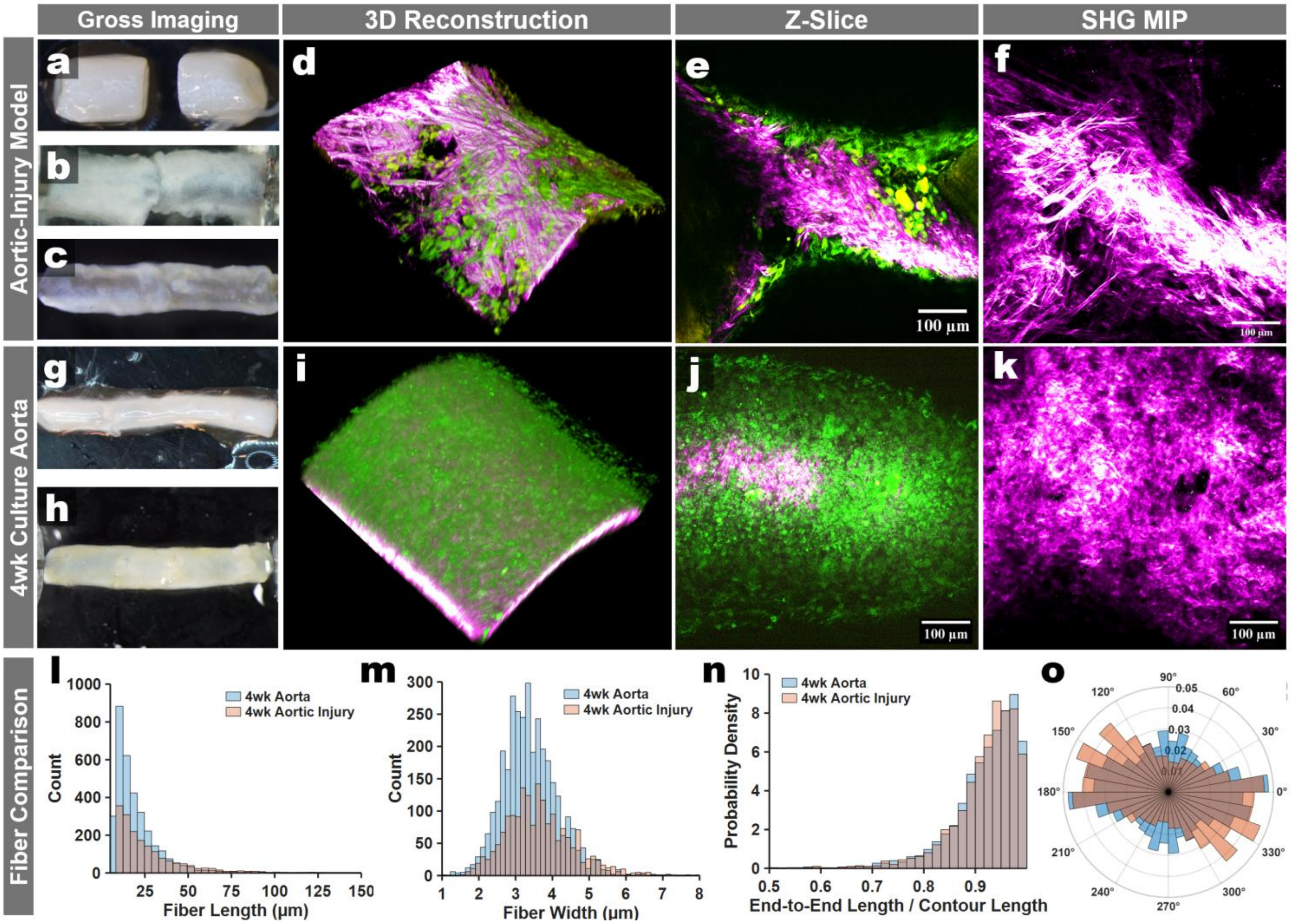


**Extended Data Fig. 1 | Label-free Multiphoton Imaging of an *Ex Vivo* Aortic Injury Model. a-c**. Gross images of the aortic injury model construct during preparation and after culture (artery–artery repair configuration). **d.** 3D volume rendering of the injury-site region showing SHG collagen signal (magenta hot) overlaid with cell-associated 2PEF as green. **e.** Representative single optical z-slice through the injury region. **f.** SHG maximum-intensity projection (MIP) highlighting fibrillar collagen organization at the injury interface. **g-h**. Gross images of a healthy (uninjured) aorta maintained in 4-week static culture as a reference tissue. **i.** 3D volume rendering of the cultured healthy aorta (SHG, magenta hot and 2PEF as green). **j**. Representative z-slice. **k.** SHG MIP of fibrillar collagen structure in the 4-week cultured healthy aorta. **l-n**. SHG fiber metrics quantified and overlaid from the 4wk cultured aorta (blue) vs. a 4wk cultured aortic injury model (transected and rejoined artery, red coloration) using established SHG fiber-extraction analysis (e.g., CT-FIRE): (**l**) fiber length distribution, (**m**) fiber width distribution, and (**n**) fiber orientation distribution (polar histogram). Scale bars, 100 µm (**e, f, j, k**).

#### Extended Data Figure 2:

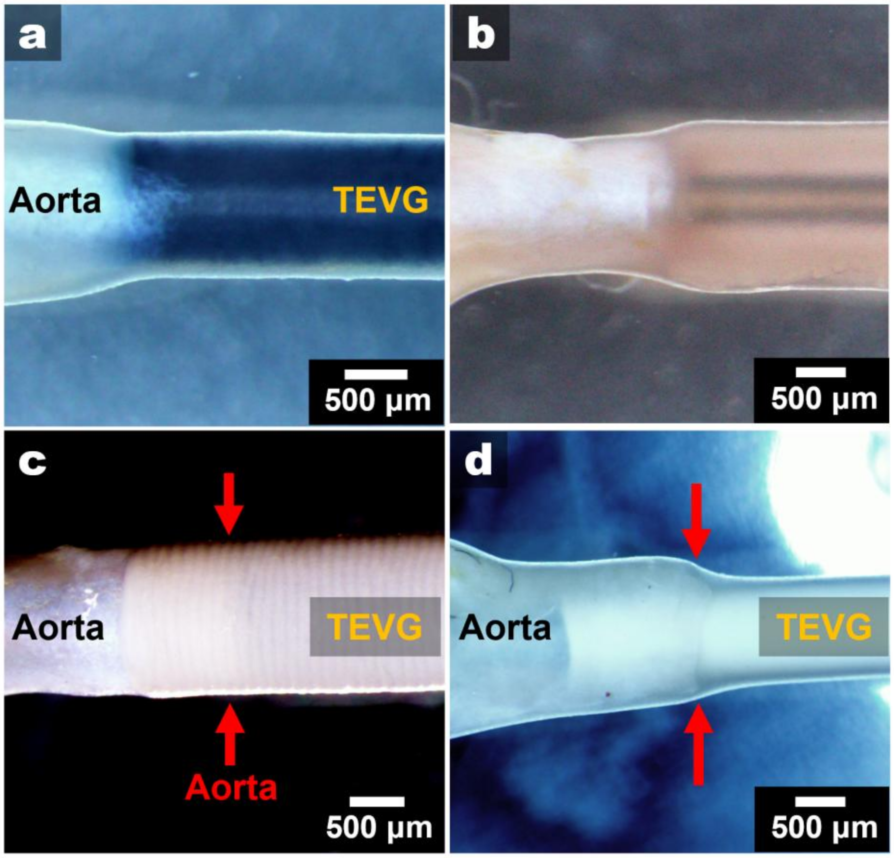


**Extended Data Fig. 2 | 3D Culture Aorta-Graft Dimensional Matching.** Gross images from various 3D cultures with different aorta-TEVG sizes. **a.** Demonstration of a dimensionally well-matched TEVG and aorta in culture, allowing for monitoring of the graft without ingrowth/outgrowth confounders. **b.** An example of a TEVG that is mismatched, but within tolerance, allowing for multiphoton imaging of cellularization and remodeling on the graft's outer surface. **c**. Depiction of a TEVG used in the culture with a significantly larger outer diameter, causing the aorta to grow on the graft's exterior. **d.** A vascular graft that is too small in outer diameter compared to the aorta. All grafts depicted are at 4-8 weeks in culture. Red arrows are pointing to the areas of ingrowth/outgrowth when mismatched in size by more than 30%.

### Extended Data Figure 3

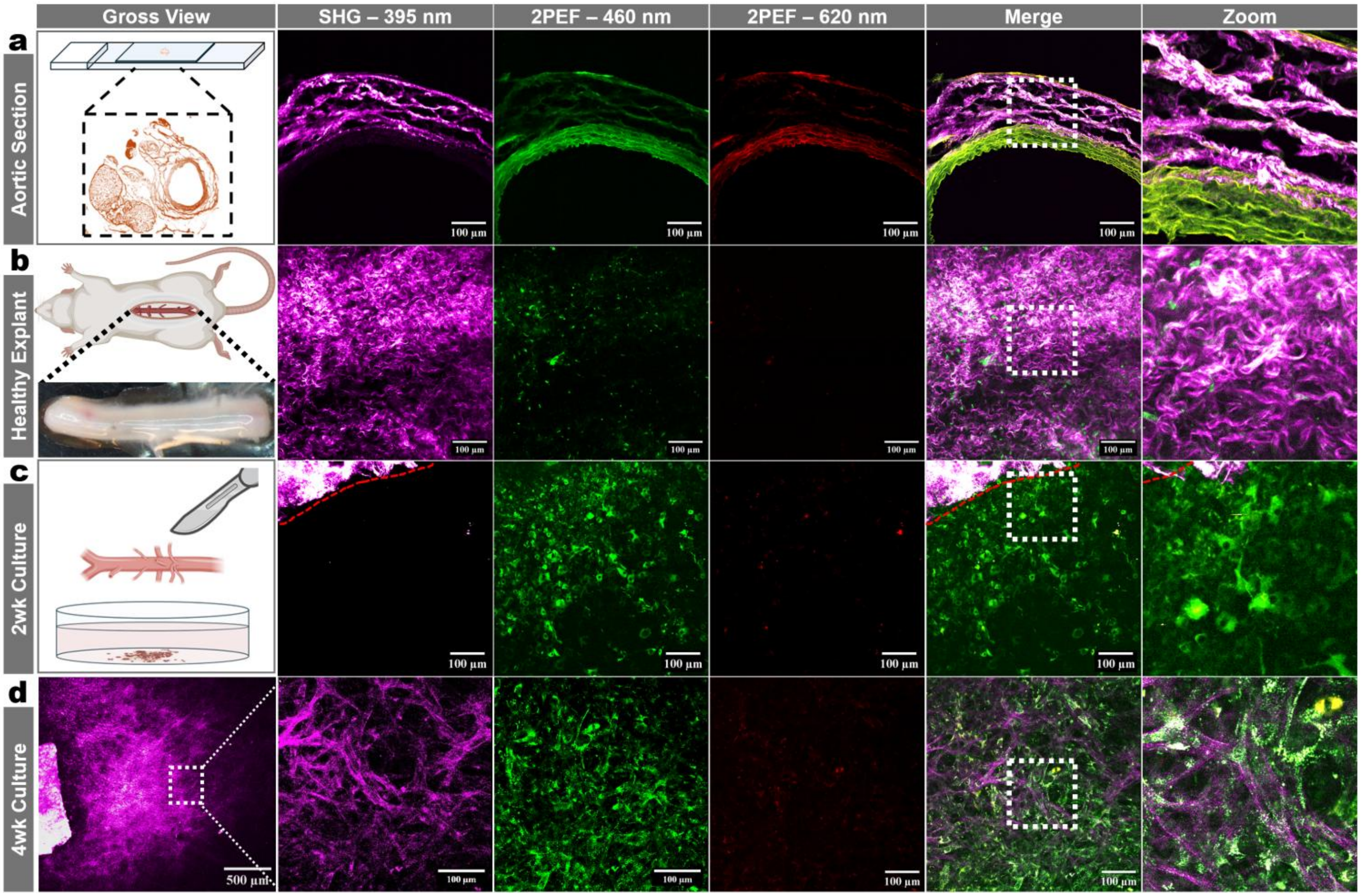


**Extended Data Fig. 3 | Multiphoton Imaging of Aortic Sections & Diced Cultures. a.** An abdominal aortic explant section from a male rat imaged under a multiphoton microscope. **b.** Live multiphoton 3D stack images of an explanted rat aorta. **c.** A live multiphoton image of an early rat aortic explant that was diced and cultured in smooth muscle cell media (SmGM-2) for 2-weeks. Top left of image shows a piece of the cut portion of aorta in view. **d.** A live multiphoton image taken from a longer-term aortic explant culture (4wks) showing high levels of cellular-associated 2PEF and collagen-associated SHG. Samples were excited at ~800 nm laser wavelength with collection across three channels simultaneously: 395/25 nm (SHG), 460/40 nm (cell-associated 2PEF), and 620/60 nm (broad 2PEF). Scale bars are 100 µm.

### Extended Data Figure 4:

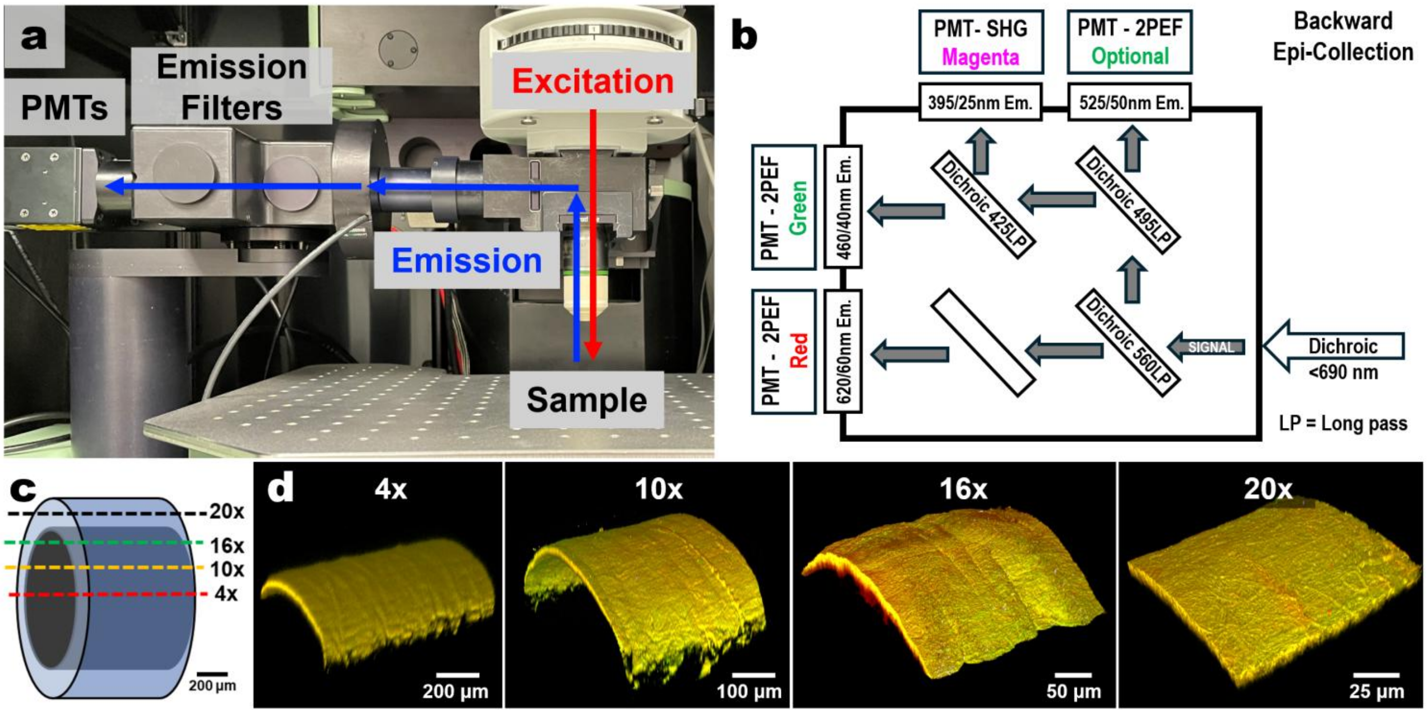


**Extended Data Fig. 4 | Multiphoton Optical Configuration and Objective Lens Depths. a.** Depiction of the gross imaging set up and light paths. **b.** A schematic of dichroic filters used for signal acquisition. The system is designed for epi (backwards) collection of SHG and 2PEF due to 3D sample thicknesses. **c.** Graphical schematic of the relative depth penetration depths when using live acquisition settings. **d.** Representative 3D reconstruction images from 3D stack imaging of the 'scaffold-only'.

### Extended Data Figure 5:

#### The platform supports multiple scaffold biomaterial choices

To assess broader applicability beyond our trilayered TEVGs, we utilized our platform to investigate electrospun grafts made from biomaterials commonly used in vascular research. As before, we assembled culture constructs and combined them with alternative biomaterials (gelatin, UBM hydrogel, PEUU, and PESBUU-50) and imaged them under matched conditions and acquisition settings for 4 weeks in reference to our original trilayered designs (**Extended Fig. 5a**). We found that electrospun porcine-derived gelatin grafts (crosslinked with genipin) exhibited high levels of cellular 2PEF and little to no SHG by 4 weeks. Notably, gelatin grafts required a 4$^{th}$ collection channel (525/50 nm) to distinguish the graft from the cellular 2PEF signatures (**Extended Fig. 5b**). We found that electrospun PESBUU-50, a zwitterionic sulfobetaine-modified PEUU, exhibited high levels of cellular 2PEF and little to no SHG signatures by 4 weeks (**Extended Data Fig. 5c**). We found that other natural biomaterials, such as decellularized porcine urinary bladder matrix (UBM), may obscure tracking of *de novo* collagen remodeling. Using established protocols, we prepared and coated UBM hydrogels upon the outer surface of a rolled spin-casted sheet of PCL and cultured the constructs as before for 4 weeks. In agreement with prior reports, the UBM-hydrogels retained SHG signal in disorganized bundle-like shapes, suggesting that the use of ECM products for SHG tracking in our platform may require alternative preparation protocols[1]. While we could not confidently distinguish de novo SHG signals, we found that the UBM-hydrogel cultures exhibited strong cellular 2PEF. In contrast, we found that electrospun PEUU grafts, which lack sulfobetaine content, exhibited robust increases in SHG and relatively low levels of cellular 2PEF (**Extended Data Fig. 5d**). We then evaluated electrospun PEUU grafts, which showed high SHG deposition and electrospun PCL, a polymer known to lack 2PEF signatures at ~800 nm excitation and found that it lacked TEVG-specific 2PEF, had minimal cellular 2PEF and that the SHG signatures could still be monitored along the graft's surfaces. PCL TEVGs had substantially lower levels of cellular 2PEF, but very high SHG levels (**Extended Data Fig. 5e,f**).

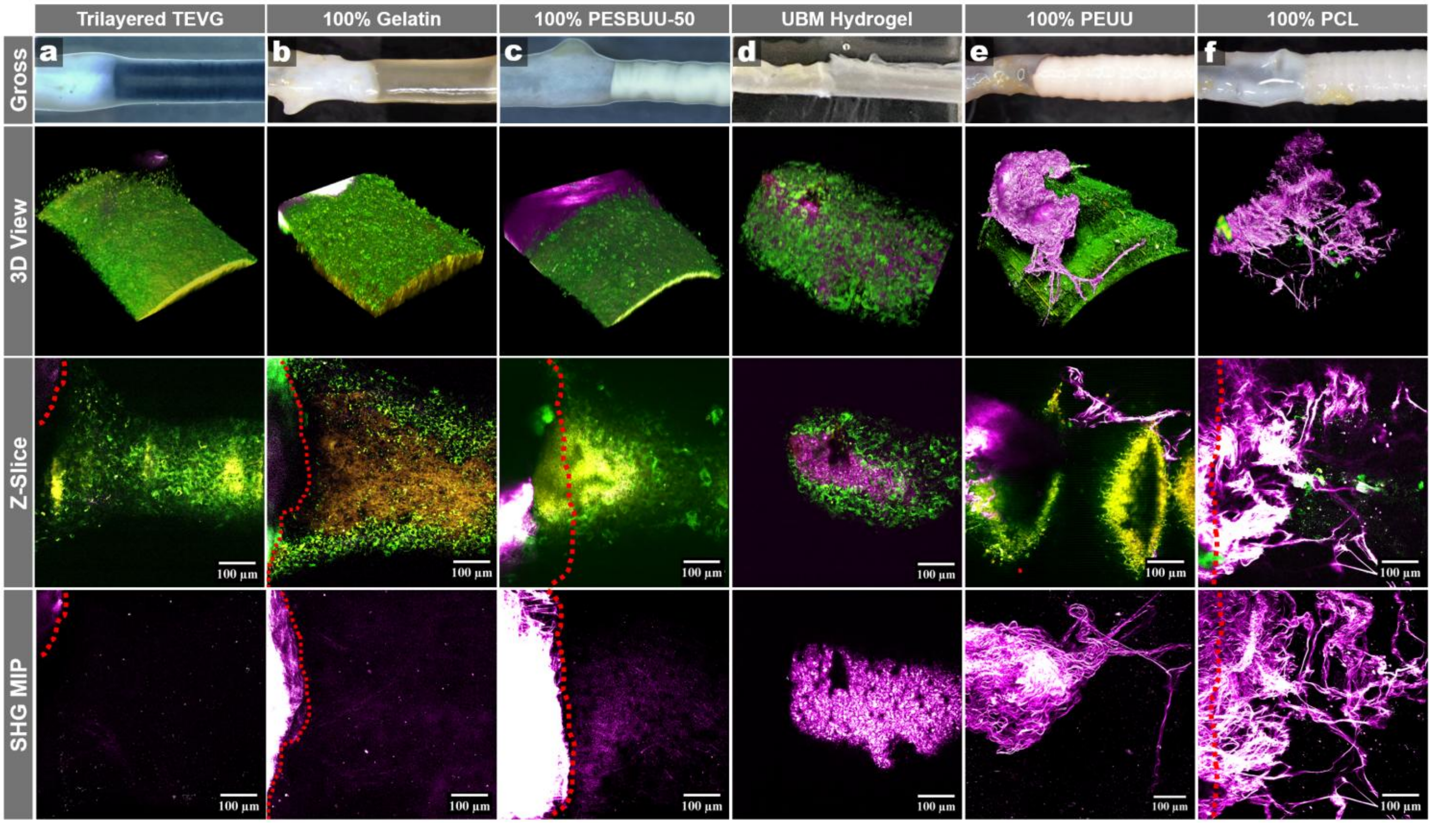


**Extended Data Fig. 5 | 3D Culture Platform Compatibility with Common TEVG Biomaterials. a-f.** Representative longitudinal live multiphoton imaging of graft interfaces in the 3D *ex vivo* culture at the 4wk timepoint. Images are displayed as merged channels under 792 nm excitation, showing SHG (magenta), cellular 2PEF (green), and TEVG (when applicable, yellow/orange). 3D reconstruction images (top), representative z-slice (middle), and SHG MIP (bottom). **a.** Images from the previous 4wk cultured trilayered TEVG, as a reference. **b.** An electrospun gelatin scaffold (crosslinked with 0.5% genipin for 24h in EtOH). 460/40 nm (orange) to identify the graft and overlap between 525/50 nm and 620/60 nm for cellular 2PEF (green). **c.** An electrospun PESBUU-50 TEVG cultured for 4wks. **d.** A spin casted PCL sheet (~25 µm in thickness) coated with a UBM hydrogel and wrapped into a TEVG cylinder at 4wks. **e.** Images from an electrospun PEUU TEVG at 4wks. **f.** An electrospun polycaprolactone (PCL) TEVG. Red dashed lines (bottom row) indicate areas where the aorta is included in the view. Scale bars are 100 µm.

#### Extended Data Figure 6:

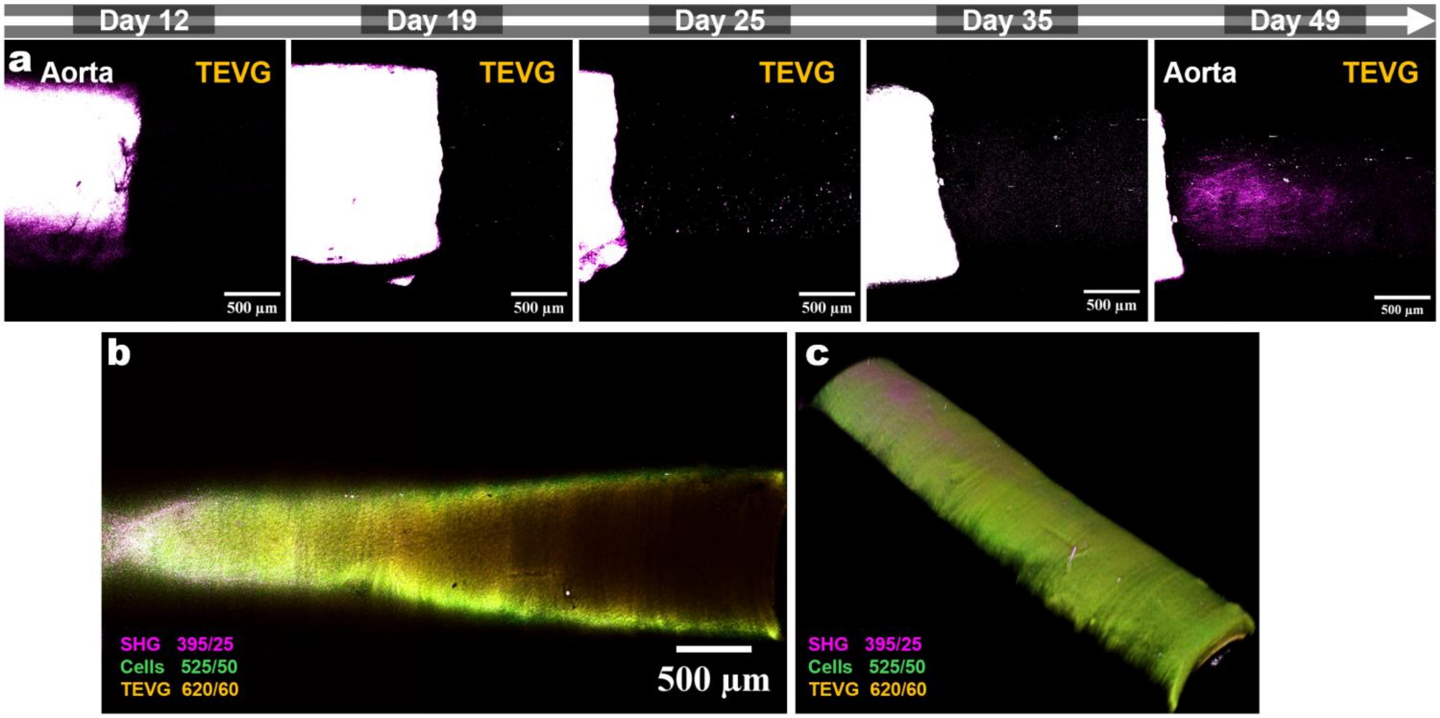


**Extended Data Fig. 6 | Tracking SHG with a 4x Objective Lens. a.** Representative images SHG (only) across time using a low-NA and low-resolution objective lens (4x). **b**. A representative z-slice image from a z-stack showing all three signals using a 4x objective. **c.** 3D reconstructions of a mosaic of 3D stack images taken of an 8wk cultured sample (single cannulation to 1 aorta) at 4x. (**b**) and (**c**) panel include all 3 channels as noted on the images.

### Extended Data Figure 7:

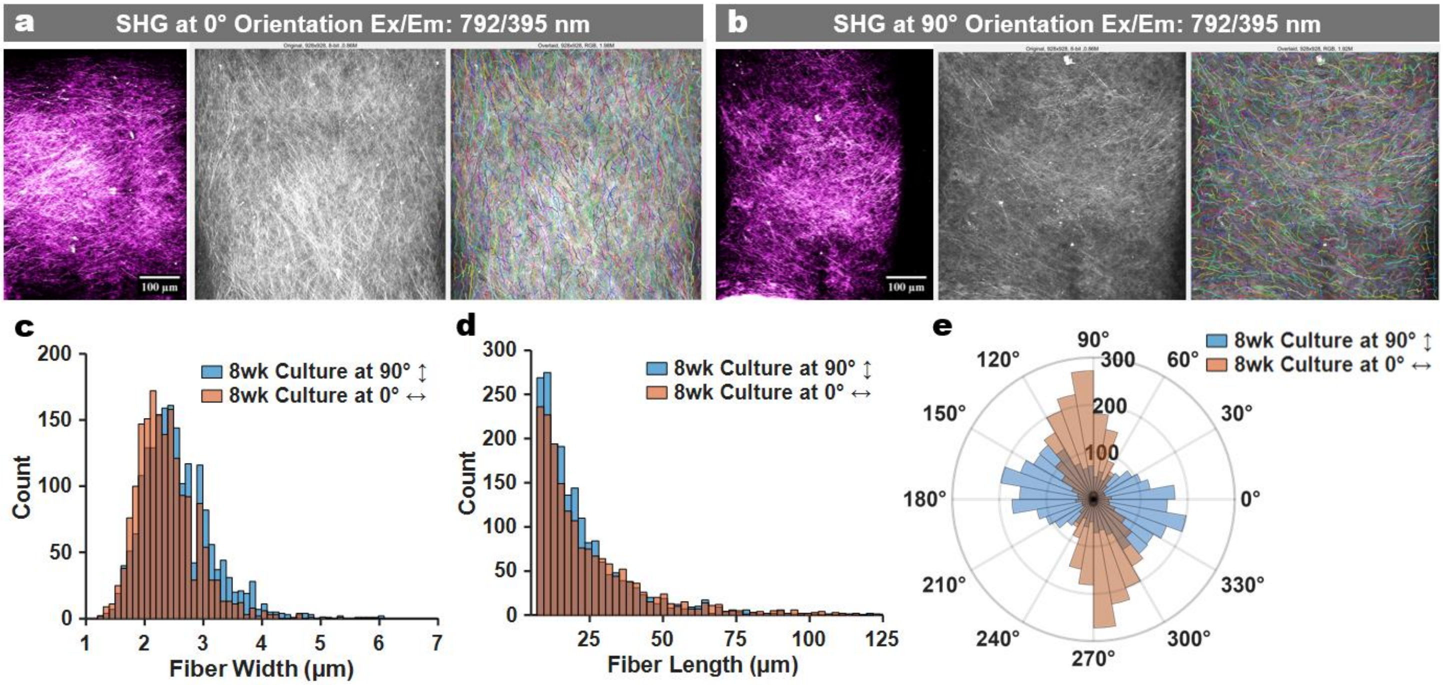


**Extended Data Fig. 7 | Sample Orientation Impacts SHG Outcomes. a.** Representative SHG channel MIP (Ex/Em:792) (left) with a CT-FIRE overlay of determined fibers (right). **b.** The same sample turned 90 degrees and imaged with the same acquisition settings (noticeably dimmer signal). **c-e.** CT-FIRE analyses of both orientations overlaid for fiber width (**c**), fiber length (**d**), and polar alignment (**e**).

#### Extended Data Figure 8:

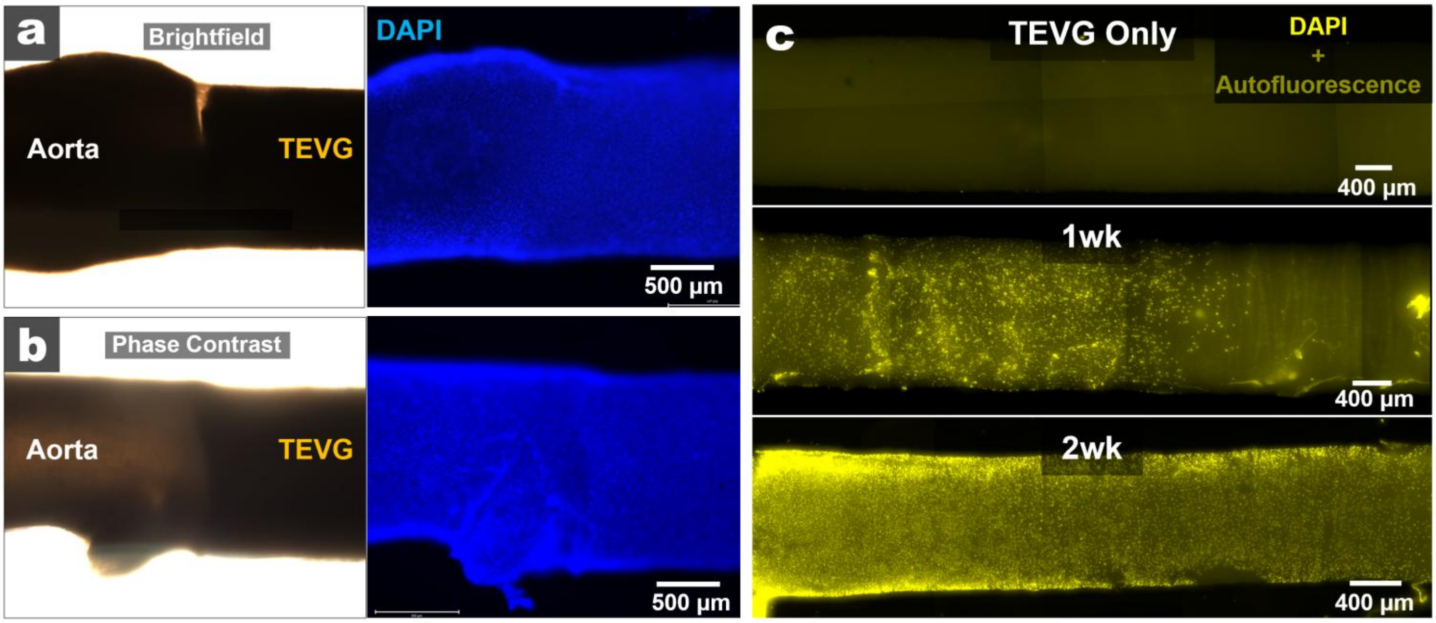


**Extended Data Fig. 8 | Fixed Sample 2PEF Validation with DAPI Staining. a.** Image of a fixed culture sample (2wks) under bright field lighting (left) and DAPI staining (right). **b.** Image of a fixed culture sample (2wks) under phase contrast imaging (left) and DAPI staining (right). **c.** Images of trilayered TEVGs after 1wk and 2wks in culture in comparison to a TEVG that cultured and DAPI stained without an aorta (top).

### Extended Data Figure 9:

#### 3D culture compatibility with fluorescent reporter strains

To expand the utility of the platform to murine models, we evaluated mouse aortic explants and a dual reporter strain in the 3D culture. We generated a tamoxifen-inducible dual fluorescent reporter to distinguish smooth muscle lineage cells (GFP) from surrounding stromal populations (tdTomato Red) for live imaging by crossing Myh11-CreER$^{T2}$ mice with the mT/mG Cre reporter line. Widefield imaging confirmed reporter signal at both the gross tissue level and in primary murine aortic cells in 2D culture, with minimal background in wild-type controls (**Extended Data Fig. 9a,b**). We found that optical clearing enabled more depth in imaging with widefield fluorescent imaging of explanted aortas, but required fixation, dehydration, and clearing protocols (**Extended Data Fig. 9c, top**). We then used multiphoton imaging to find that the tdTomato and GFP signatures were weak under our standard live imaging acquisition settings, and that the expected 2-photon excitation of 900 nm was sufficient to visualize the GFP and tdTomato red signal within live aortas (**Extended Data Fig. 9d, bottom**). We then verified that the dual reporter aortas could successfully be integrated into the 3D culture model and be imaged using our live acquisition settings to see cells, SHG, and TEVG, as well as utilize different excitations to visualize the GFP$^+$ smooth muscle cells (**Extended Fig 9e, top and bottom panels, respectively**).

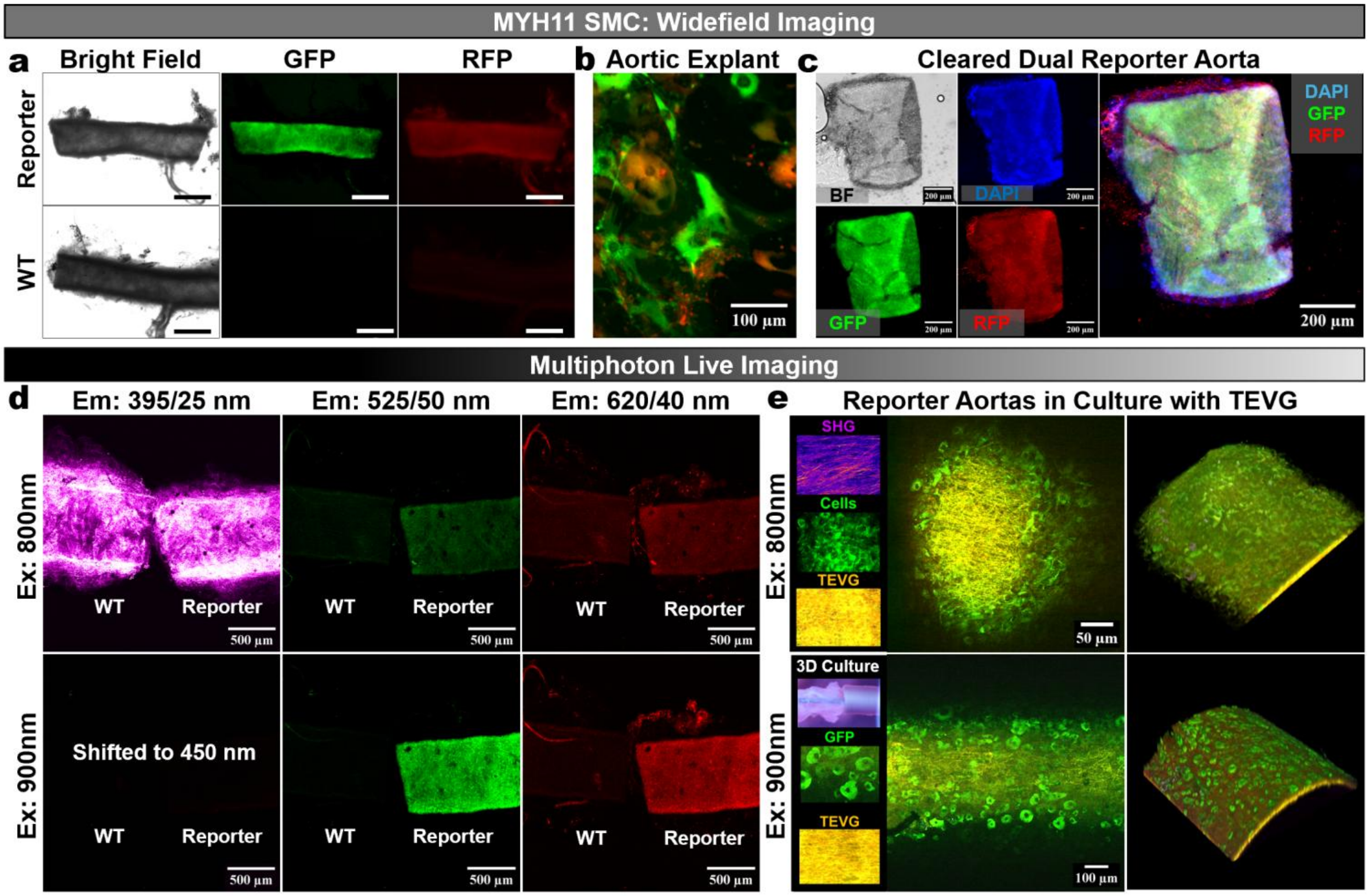


**Extended Data Fig. 9 | Murine & Reporter-Strain Compatibility in the 3D Culture Platform. a**. Gross validation of a dual-reporter mouse line following tamoxifen induction of Cre recombination, with ubiquitous tdTomato (RFP) labeling of all cells and GFP labeling of MYH11 expressing smooth muscle cells (SMCs). Representative brightfield, GFP, and RFP fluorescence images compare reporter aortic tissue to wild-type (WT) controls to confirm reporter specificity at the tissue scale. **b.** Higher-magnification fluorescence image of primary murine aortic cells in 2D culture demonstrating cellular-level reporter expression and co-localization patterns consistent with the dual-reporter design. **c**. Terminal optical clearing enables limited volumetric imaging of the dual reporter signal under widefield fluorescence, shown as brightfield (BF), DAPI nuclear stain, GFP, RFP, and a merged reconstruction (DAPI/GFP/RFP). **d.** 4x Live multiphoton imaging of WT and reporter aortas to map signatures across the standardized collection channels. Under ~800 nm excitation (top) and ~900 nm (bottom), signal is shown for SHG (395/25 nm; magenta), the GFP-compatible window (525/50 nm; green), and the tdTomato/RFP-compatible window (620/60 nm; red). **e, top.** 16x representative z-plane image (left) and 3D reconstruction under ~800 nm excitation showing cellular 2PEF and scaffold 2PEF. **e**, **bottom**. Increasing the excitation wavelength to ~900 nm enhances visualization of the GFP reporter, with corresponding channel readouts shown for comparison (16x). Scale bars in (a) are 1 mm, (c) 200 µm, (d) 500 µm, (e, top) 50 µm, & (e, bottom) 100 µm.

### Extended Data Figure 10:

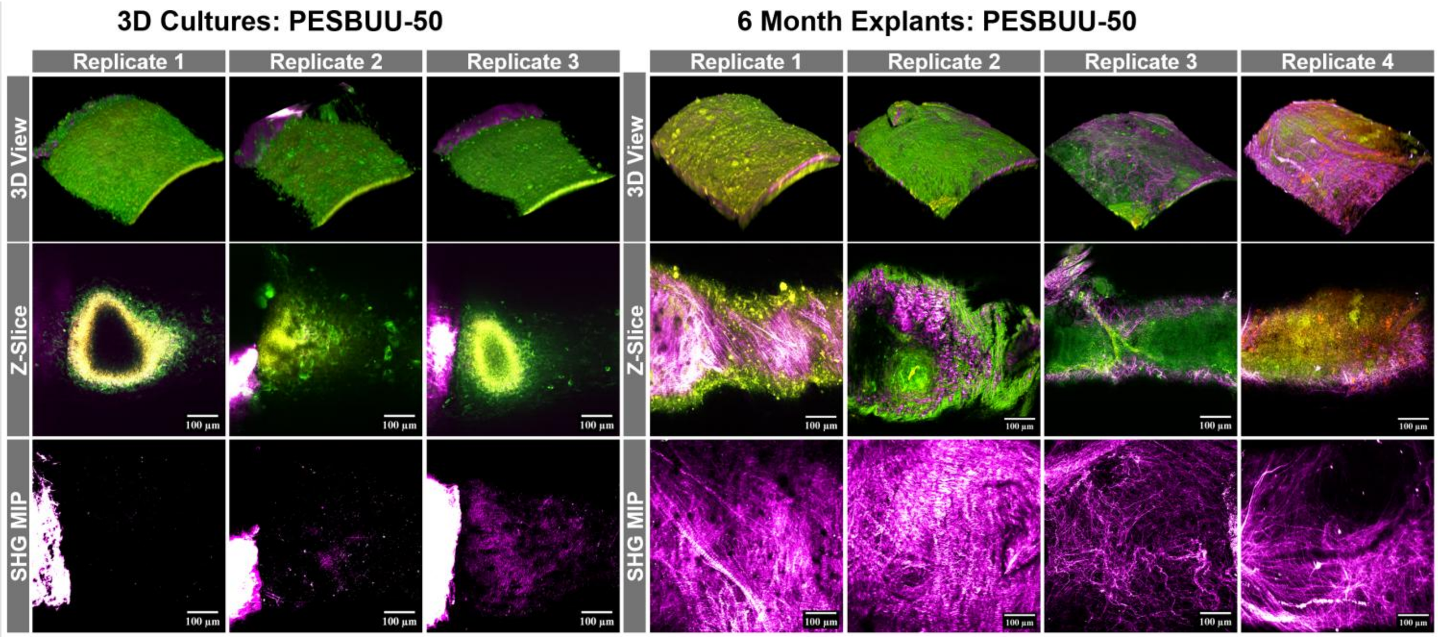


**Extended Data Fig. 10 | PESBUU-50 6-Month Explants vs 8-Week Cultures.** Replicate multiphoton images shown for PESBUU-50 grafts explanted after 6-months as interpositional grafts in comparison to PESBUU-50 replicates from the 3D cultures platform after 8wks. For space, the 4$^{th}$ replicate for the PESBUU-50 culture is shown in main figures. Scale bars are 100 µm.

### Extended Data Table 1: Reagents & Supplies

| Vascular Graft Fabrication Supplies | Catalog # / ID | Vendor | Supplier (If Applicable) |
|---|---|---|---|
| 1,1,1,3,3,3-Hexafluoro-2-propanol (HFIP) | H1008 | Spectrum Chemical | - |
| Type A Gelatin from Porcine Skin | G1890 | MilliporeSigma | - |
| Polycaprolactone 80kDa | 440744 | MilliporeSigma | - |
| Genipin | 6902-77-8 | FujiFilm | Fisher Scientific |
| PEUU | - | - | In-house preparation |
| PESBUU-50 | - | - | In-house preparation |
| **3D Culture Supplies** | **Catalog # / ID** | **Vendor** | **Supplier (If Applicable)** |
| SmGm2 Media with Bullet Kit | CC-3182 | Lonza | - |
| PTFE coated mandrels (<0.5mm OD) | 206033 | Applied Plastics | - |
| Polycarbonate filament (Ultimaker) | 1640 | Ultimaker | Dynamism |
| 1x Sterile DPBS | 14190235 | Gibco | Thermo Fisher Scientific |
| Pen/Step 10,000 U | 15140122 | Gibco | Thermo Fisher Scientific |
| Live Cell Imaging Solution (optional) | A59688DJ | Gibco | Thermo Fisher Scientific |
| FluoroBrite DMEM | A1896701 | Gibco | Thermo Fisher Scientific |
| Amphotericin B | 15290018 | Gibco | Thermo Fisher Scientific |
| 100mm x 25mm Petri Dishes | FB0875711 | FisherBrand | Fisher Scientific |
| 60mm Petri Dishes | 50828744 | CELLTREAT | Fisher Scientific |
| 35mm Petri Dishes | 627161 | Greiner Bio-One | Thermo Fisher Scientific |
| 6-well Plates | 140675 | Thermo Scientific Nunc | Thermo Fisher Scientific |
| SylGard 184 Kit (optional PDMS) | 4019862 | Dow | Krayden |
| Whatman lens cleaning tissue, Grd. 105 | WHA2105841 | Whatman | MilliporeSigma |
| **Proteins for 3D Culture** | **Catalog # / ID** | **Vendor** | **Supplier (If Applicable)** |
| rhTGFb1 carrier-free | 240B010/CF | R&D Systems | - |
| rhTGFb2 carrier-free | 302-B2-010/CF | R&D Systems | - |
| rhTGFb3 carrier-free | 243B3010/CF | R&D Systems | - |
| **qRT-PCR Gene Expression** | **Catalog # / ID** | **Vendor** | **Supplier (If Applicable)** |
| RNALater | AM7020 | Ambion | Thermo Fisher Scientific |
| Trizol Reagent | 15596018 | Invitrogen | Thermo Fisher Scientific |
| Biomasher 2 (pestles+tubes) | K7496250030 | Nippi | Fisher Scientific |
| RNA suitable Chloroform | C2432-4X25ML | MilliporeSigma | - |
| RNEasy Microkit PLUS | 74034 | Qiagen | - |
| TaqMan Master Mix | 4370048 | Applied Biosystems | Thermo Fisher Scientific |
| 0.2mL PCR Tube Strips | 14230215 | FisherBrand | Fisher Scientific |
| SuperScript IV VILO | 11756050 | Invitrogen | Thermo Fisher Scientific |
| DNA LoBind 0.5mL tubes | 05-414-202 | Eppendorf | Fisher Scientific |
| Optical Film | 4311971 | Applied Biosystems | Thermo Fisher Scientific |
| Optical Plates | N8010560 | Applied Biosystems | Thermo Fisher Scientific |
| TaqMan Probe: *Rer1* (MGB-FAM) | Rn01516516_m1 | Applied Biosystems | Thermo Fisher Scientific |
| *TaqMan Probe: Hprt1* (MGB-FAM) | Rn01527840_m1 | Applied Biosystems | Thermo Fisher Scientific |
| *TaqMan Probe: Gapdh* (MGB-FAM) | Rn01775763_g1 | Applied Biosystems | Thermo Fisher Scientific |
| *TaqMan Probe: Acta2* (MGB-FAM) | Rn01759928_g1 | Applied Biosystems | Thermo Fisher Scientific |
| *TaqMan Probe: Myh11* (MGB-FAM) | Rn01530321_m1 | Applied Biosystems | Thermo Fisher Scientific |
| *TaqMan Probe: Lox* (MGB-FAM) | Rn01491829_m1 | Applied Biosystems | Thermo Fisher Scientific |
| *TaqMan Probe: Col1a1* (MGB-FAM) | Rn01463848_m1 | Applied Biosystems | Thermo Fisher Scientific |
| *TaqMan Probe: Col3a1* (MGB-FAM) | Rn01437681_m1 | Applied Biosystems | Thermo Fisher Scientific |
| **Histological Staining** | **Catalog # / ID** | **Vendor** | **Supplier (If Applicable)** |
| Rabbit Anti-Rat-Vimentin | ab92547 | Abcam | - |
| Rabbit Anti-Rat-MYHC11 | ab224804 | Abcam | - |
| Rabbit Anti-Rat α-Smooth Muscle Actin | ab7817 | Abcam | - |
| Goat anti-Rabbit IgG (AF488) | ab150077 | Abcam | - |
| Picrosirius Red Stain (Connective Tissue) | ab150681 | Abcam | - |
| **Immunofluorescent Staining/Dyes** | **Catalog # / ID** | **Vendor** | **Supplier (If Applicable)** |
| NucBlue Live ReadyProbes Hoechst 33342 | R37605 | Invitrogen | Thermo Fisher Scientific |
| NucBlue Fixed Cell ReadyProbes DAPI | R37606 | Invitrogen | Thermo Fisher Scientific |

| Extended Data Table 2: Electrospinning Parameters for Trilayered TEVGs | | | |
|---|---|---|---|
| **Layer** | **Lumen** | **Media** | **Outer** |
| **Polymer** | PESBUU-50 | PEUU:Gelatin | PEUU:Gelatin |
| **Polymer Ratio** | 100 % | 80:20 | 20:80 |
| **Conc. (w/v %)** | 8 % in HFIP | 10 % in HFIP | 10 % in HFIP |
| **Dispensing Capillary ID** | 0.81 mm (18G) Stainless Steel | 0.62 mm (20G) Stainless Steel | 0.62 mm (20G) Stainless Steel |
| **Dispense Volume Dispense Rate** | 30 µL 10 µL/min | 65 µL 10 µL/min | 125 µL 10 µL/min |
| **Nozzle Translation Distance & Rate** | 80 mm 30 mm/s | 80 mm 30 mm/s | 80 mm 30 mm/s |
| **Mandrel OD** | 1.1 mm | 1.1 mm | 1.1 mm |
| **Mandrel Length** | 150 mm | 150 mm | 150 mm |
| **Mandrel Coating** | PTFE | PTFE | PTFE |
| **Mandrel Rotation** | 1000 RPM | 1000 RPM | 1000 RPM |
| **Working Distance** | 12 cm | 10 cm | 10 cm |
| **Voltage Settings** | +20 kV Nozzle -4 kV Mandrel | +20 kV Nozzle -4 kV Mandrel | +20 kV Nozzle -4 kV Mandrel |
| **Temp. & Humidity** | 21°C, 30 % | 21°C, 30 % | 21°C, 40 % |
| **Crosslinking** | 24hrs in 0.5% Genipin in 200 proof EtOH at 37°C | | |
| **Portion Used** | Central 50 mm (±25 mm from center) | | |

| Extended Data Table 3: Tubular Biaxial Testing | |
|---|---|
| **Submerged in** | DPBS or Krebs-Henseleit |
| **Temperature** | 37.0 ± 2.0 °C |
| **Initial Axial Load** | Taut (~0.010 N) |
| **Initial Luminal Pressure** | 0.1 ± 0.05 mmHg |
| **Loading Scheme** | Cyclic, Sinusiodal |
| **Loading Rate(s)** | 1 mmHg/s; 0.5mm/s |
| **Cyclic Applied Pressure** | 0.1 to 120.0 ± 1 mmHg |
| **Sampling Rate** | 5 Hz |
| **No. of PreCond. Cycles** | 9 |
| **Controller Type** | PID |
| **No. of Diameter ROIs** | 6 pairs |
| **Axial Length of ROI** | 5 mm |

$$\textit{Compliance} = \frac{\left(\frac{OD_{120} - OD_{70}}{OD_{70}}\right)}{(120 - 70)\ \text{mmHg}}$$

| Extended Data Table 4: Multiphoton Hardware, Optical Filters, & PMTs | | | |
|---|---|---|---|
| **Multiphoton Component** | **Type** | **Vendor** | **Catalog #** |
| TriM Scope II | Upright | LaVision Biotec | - |
| InSight Deep See + Dual laser | | Newport | InSight DS+ |
| **Photomultiplier Tube (PMT)** | **Type** | **Vendor** | **Catalog #** |
| Non-descanned PMT 1 | GaAsP | Hamamatsu | H7422-40-LV 5M |
| Non-descanned PMT 2 | GaAsP | Hamamatsu | H7422-40-LV 5M |
| Non-descanned PMT 3 | GaAsP | Hamamatsu | H7422-40-LV 5M |
| Non-descanned PMT 4 | Bialkali | Hamamatsu | H6780-01-LV 5M |
| **Objectives** | **NA** | **Vendor** | **Catalog #** |
| Olympus XL Fluor 2x | 0.14 | Olympus | AMEP4751 |
| Olympus XL Fluor 4x | 0.28 | Olympus | AMEP4979 |
| UM Plan Fl N 10x | 0.30 | Olympus | 1-U2M583 |
| Nikon LWD 16x | 0.80 | Nikon | CFI75 |
| W Plan-Apochromat 20x | 1.00 | Zeiss | 421452-9980-000 |
| **Filter** | **Type** | **Vendor** | **Catalog #** |
| 377/50 nm | Em | Chroma Technology | ET377/50m |
| 395/25 nm | Em | Chroma Technology | ET395/25x |
| 460/40 nm | Em | Chroma Technology | ET460/40m |
| 525/50 nm | Em | Chroma Technology | ET525/50m |
| 620/60 nm | Em | Chroma Technology | ET620/60m |
| 425LP | DLP | Chroma Technology | T425lpxr |
| 495LP | DLP | Chroma Technology | T495lpxr |
| 560LP | DLP | Chroma Technology | T560lpxr |
| 690LP | DLP | Chroma Technology | T690lpxr |
| DLP/LP = Dichroic long pass, Em = Emission (bandpass), PMT = Photomultiplier tube | | | |

| Extended Data Table 5: Live Imaging Settings 16x | |
|---|---|
| **Excitation** | 792 nm |
| **Laser Power** | 13.5mW |
| **Time per Z-Slice** | 22,265 ms |
| **Pixel Dwell Time** | 4.4 µs |
| **Scan Frequency** | 200 Hz |
| **Line Averages** | 4 |
| **Avg. Z-Depth** | 200 µm |
| **Z-Step Size** | 1.5 - 2.0 µm |

| Extended Data Table 6: EVOS M7000 Optical Filters and Objectives | | |
|---|---|---|
| **Filter Cubes** | **Excitation/Emission** | **Catalog Number** |
| DAPI | Ex: 357/44 Em: 447/60 | AMEP4950 |
| GFP | Ex: 482/25 Em: 524/24 | AMEP4951 |
| Texas Red | Ex: 585/29 Em: 628/32 | AMEP4955 |
| CY5 | Ex: 635/18 Em: 692/40 | AMEP4956 |
| RFP | Ex: 542/20 Em: 593/40 | AMEP4952 |
| CFP | Ex: 445/45 Em: 510/42 | AMEP4953 |
| YFP | Ex: 500/24 Em: 542/27 | AMEP4954 |
| CY7 | Ex: 716/40 Em: 794/32 | AMEP4967 |
| **Objective Lens** | **EVOS M7000 Objective Lens Name** | **Catalog Number** |
| 4x | OBJ FL 4X LWDPH 0.13NA/10.58WD | AMEP4980 |
| 10x | OBJ FL 10X LWDPH 0.30NA/7.13WD | AMEP4981 |
| 20x | OBJ FL 20X LWDPH 0.45NA/6.12WD | AMEP4982 |
| 40x | OBJ OLY FL 40X 0.6/2.7-4.4 | AMEP4764EO |
| 60x | OBJ OLY FL 60X 0.9NA/0.2WD | AMEP4849 |

| Extended Data Table 7: Electrospinning Parameters for Single Layer TEVGs | | | | |
|---|---|---|---|---|
| **Alternative TEVGs** | | | | |
| **Polymer** | **PCL** | **Gelatin** | **PEUU** | **PESBUU-50** |
| **Conc. (w/v %)** | 8 % in HFIP | 10 % in HFIP | 8 % in HFIP | 8 % in HFIP |
| **Dispensing Capillary ID** | 125 µL 15 µL/min | 125 µL 10 µL/min | 350 µL 10 µL/min | 350µL 10 µL/min |
| **Dispense Volume Dispense Rate** | 0.62 mm (20G) Stainless Steel | 0.62 mm (20G) Stainless Steel | 0.62 mm (20G) Stainless Steel | 0.81 mm (18G) Stainless Steel |
| **Nozzle Translation Distance & Rate** | 80 mm 30 mm/s | 80 mm 30 mm/s | 80 mm 30 mm/s | 80 mm 30 mm/s |
| **Mandrel OD** | 1.1 mm | 1.1 mm | 1.1 mm | 1.1 mm |
| **Mandrel Length** | 150 mm | 150 mm | 150 mm | 150 mm |
| **Mandrel Coating** | PTFE | PTFE | PTFE | PTFE |
| **Mandrel Rotation** | 300 RPM | 300 RPM | 1000 RPM | 1000 RPM |
| **Working Distance** | 15 cm | 12 cm | 10 cm | 12 cm |
| **Voltage Settings** | +15 kV Nozzle -1 kV Mandrel | +15 kV Nozzle -1 kV Mandrel | +20 kV Nozzle -4 kV Mandrel | +20 kV Nozzle -4 kV Mandrel |
| **Temp. & Humidity** | 21°C, 30 % | 21°C, 30 % | 21°C, 30 % | 21°C, 30 % |
| **Crosslinking** | None | 0.5% Genipin 24h | None | None |
| **Portion Used** | Central 50 mm | Central 50 mm | Central 50 mm | Central 50 mm |
